# RNA Helicase A promotes small RNA biogenesis and sorting in germ cells

**DOI:** 10.1101/2025.05.11.653140

**Authors:** Olivia J. Gaylord, Jordan S. Brown, Wei-Sheng Wu, Heng-Chi Lee

## Abstract

In germ cells, small RNAs function as a defense system to silence invading RNAs like viruses and transposons to protect genome integrity. The ability of small RNAs to robustly silence diverse RNA sequences prompts the question of how endogenous mRNAs avoid this silencing. In *C. elegans*, small RNAs bound by the Argonaute CSR-1 protect endogenous mRNAs from silencing, while also fine-tuning a subset of these mRNAs. Here, we identify RNA Helicase A (RHA-1) as a key regulator of CSR-1 small RNA biogenesis and function in mRNA fine-tuning. RHA-1 localizes to germ granules dependent on EGO-1, which synthesizes CSR-1 small RNAs. We find RHA-1 promotes small RNA production from the 5’ regions of mRNAs and small RNA sorting to CSR-1. Loss of RHA-1 leads to elevated CSR-1 target mRNA levels and compromised fertility. Our study highlights the importance of small RNA regulation, mediated by RHA-1, to protect endogenous gene expression programs and germ cell function.

## Introduction

In animals, small RNAs are bound by the conserved family of Argonaute proteins to form complexes that can induce RNA silencing by binding complementary mRNA sequences leading to mRNA degradation, translational repression, or transcriptional silencing (*1*, *2*). Within germ cells, several classes of small RNAs, including micro-RNAs (miRNAs), short-interfering RNAs (siRNAs), and piwi-interacting RNAs (piRNAs) are produced by different biogenesis factors and bound by distinct Argonaute members (*3*). Together these small RNA classes perform crucial roles in genome integrity, germ cell development and differentiation, and overall germline fertility (*1*). PiRNAs and siRNAs perform well-established roles in defending the genome from mutagenic invading RNAs including viral RNAs and transposons (*4–7*). Outside of this role in genome defense, more recent work uncovered that piRNAs and endogenous siRNAs also regulate endogenous protein-coding mRNAs during sperm development in mice, fruit flies, and worms (*8–10*). These expanded gene regulatory roles complicate our understanding of how small RNA classes establish and maintain their mRNA target specificity to facilitate multiple biological processes during germ cell development.

In *C. elegans*, the most abundant class of small RNAs are endogenous siRNAs termed 22G-RNAs. This class of small RNAs is bound by a group of worm-specific Argonautes termed WAGOs and Argonaute CSR-1 (*11*). These 22G-RNAs are produced directly from mRNA transcripts that serve as templates for RNA-dependent RNA polymerases (RdRPs) EGO-1 and RRF-1 (*12–14*). The production of CSR-1 22G-RNAs requires EGO-1, whereas WAGO-1 22G-RNAs can be produced by either EGO-1 or RRF-1. WAGO-1 and CSR-1 bound 22G-RNAs target distinct sets of mRNAs and induce different gene regulatory outcomes. While WAGO-1 bound 22G-RNAs primarily function downstream of piRNAs and siRNAs to robustly silence transposons and viral RNAs, CSR-1 bound 22G-RNAs protect endogenous protein-coding mRNAs from piRNA silencing (*15–19*). Together, this system allows for robust defense against invading mutagenic RNAs, while preventing aberrant silencing of endogenous genes. Disruption to this small RNA system often results in germ cell development and differentiation defects, especially in combination with environmental stress like elevated temperature (*20*).

In addition to mRNA protection, more recent work discovered that CSR-1 also fine-tunes or decreases the mRNA levels for a subset of endogenous mRNA targets during embryo and adult germ cell development (*21–23*). This subset of mRNAs are termed CSR-1 slicer targets because they require CSR-1 slicer activity for mRNA fine-tuning. CSR-1 slicer activity performs a critical role in gene regulation as CSR-1 slicer mutants display robust defects in fertility and embryo viability nearly identical to *csr-1* knockout animals (*21*, *22*). Notably, WAGOs do not possess the critical catalytic residues for slicer activity, marking a key difference in CSR-1 and WAGO-1 gene regulatory pathways (*11*). Given that CSR-1 and WAGO-1 perform distinct regulatory roles in germ cells but share the RdRP EGO-1 for producing their associated 22G-RNAs, how 22G-RNAs are correctly sorted to CSR-1 or WAGO-1 remains an important area of investigation.

Here, we demonstrate that the conserved RNA Helicase A (RHA-1) promotes 22G-RNA biogenesis and sorting to CSR-1. Loss of RHA-1 leads to reduced CSR-1 bound 22G-RNAs and increased WAGO-1 bound 22G-RNAs derived from endogenous protein-coding mRNAs. These small RNA defects correspond to increased mRNA levels of CSR-1 target genes and spermatogenesis-enriched genes, suggesting disruption of CSR-1 target fine-tuning and germline gene regulation. Consistent with a role in CSR-1 small RNA synthesis, we find RHA-1 localizes to perinuclear germ granules within the E compartment where EGO-1 is also enriched. Moreover, RHA-1 localization to germ granules depends on both EGO-1 and CSR-1. Overall, loss of RHA-1 leads to functional defects in germ cell differentiation, fertility, and germline immortality.

## Results

### RHA-1 ATPase activity promotes RNAi and fertility in germ cells

Previous reports identified *rha-1/dhx-9*, a conserved member of the DExH family of RNA helicases, as required for various processes of genome integrity, including RNAi-mediated gene silencing in fruit flies, human cells, and worms (*24–28*). Since these reports, how RHA-1 contributes to genome integrity remains unknown. In follow up, we aimed to confirm whether RHA-1 is required for RNAi by feeding animals bacteria expressing-dsRNA against germline genes *pos-1* and *mex-3*, and somatic genes *dpy-13* and *unc-22*. For germline genes, wild type animals showed entirely unhatched embryos, indicative of successful RNAi knockdown of *pos-1* and *mex-3* genes. In contrast, *rha-1(tm329)* mutants showed nearly all hatched embryos, indicating RHA-1 is required for RNAi against germline genes (Fig. 1A). Nonetheless, both wild type and *rha-1(tm329)* mutants showed successful RNAi knockdown of somatic genes *dpy-13* and *unc-22*, based on the visible phenotypes of shortened dumpy bodies and uncoordinated twitching, respectively (Fig. 1B). Together, this data demonstrates RHA-1 is only required for RNAi function in germ cells but not somatic cells. This places RHA-1 amongst a small set of genes including CSR-1 and EGO-1 that are similarly required for RNAi only in germ cells (*12*, *29*).

**Figure 1.**
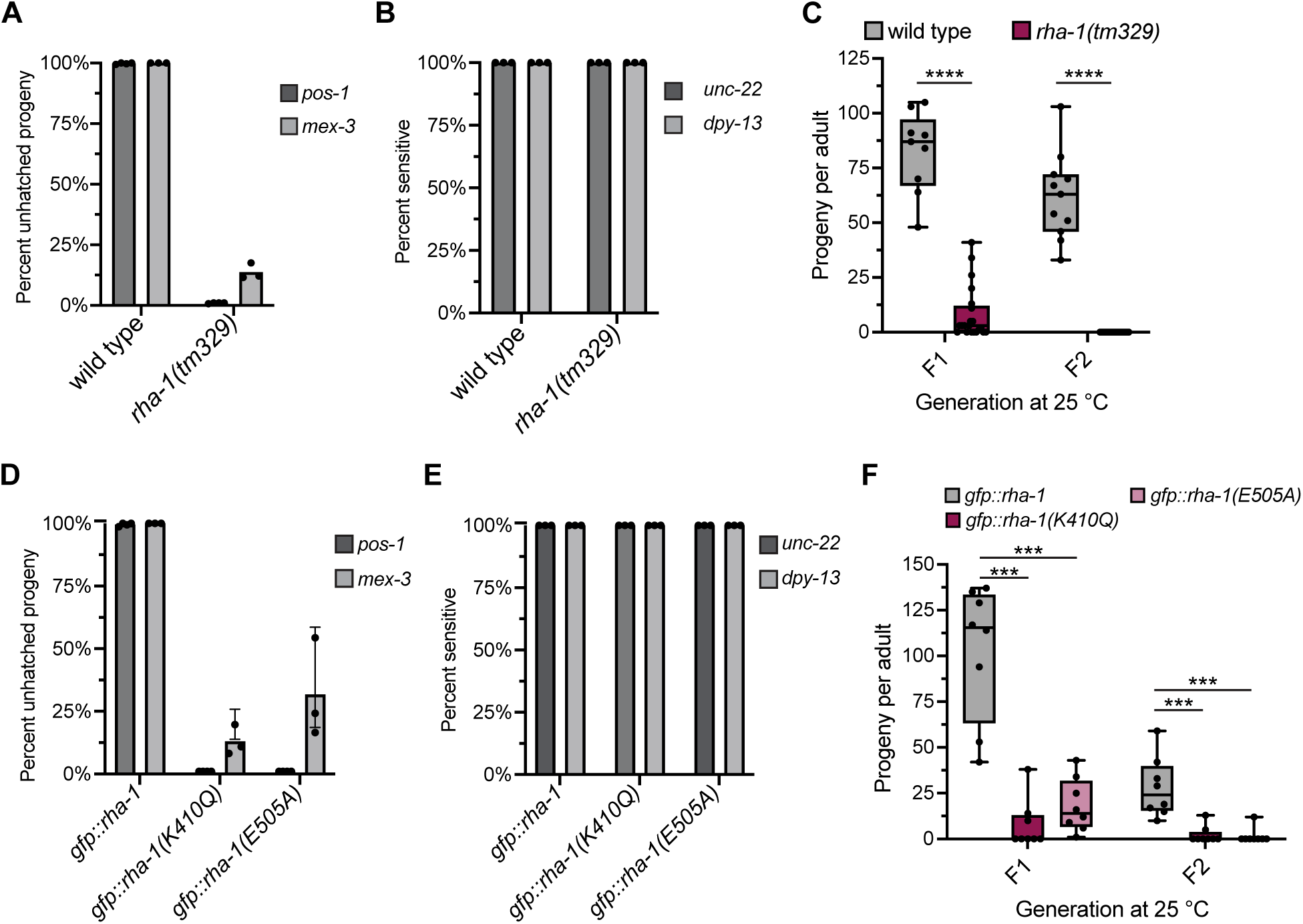
RHA-1 is required for germline RNAi and germline immortality. (**A** and **D**) Quantification of unhatched embryos of the indicated animals after feeding RNAi targeting *pos-1* and *mex-3*. Data are presented as mean values + / - SD of three biologically independent samples. (**B** and **E**) Quantification of twitching or dumpy short body phenotypes of the indicated animals after feeding RNAi targeting *unc-22* and *dpy-13* respectively. Percent sensitive indicates ratio of worms with the successful target knockdown phenotypes. Data are presented as mean values + / - SD of three biologically independent samples. (**C** and **F**) Brood sizes of the indicated animals grown at 25 °C for one (F1) or two (F2) generations. Single L3 stage animals were transferred to individual plates at each generation and number of progeny per worm was measured. *N =* 8-12 animals. For the boxplots, the line indicates the median value, the box indicates the first and third quartiles, and the whiskers indicate the min and max values. Two-tailed p values were calculated using Mann-Whitney-Wilcoxon test. ***, p<0.001; ****, p<0.0001.

In addition, these germ cell-only RNAi factors have been shown to play critical roles in germline fertility and immortality, especially at elevated growth temperature (25 °C) (*29–31*). A previous report observed reduced fertility in *rha-1(tm329)* mutants, particularly in elevated temperature (*24*). However, it was later discovered an unlinked mutation in the gene *haf-6* in the strain background may have increased the penetrance of the reported phenotypes. After outcrossing the *haf-6* mutation from the background, we examined the fertility of the *rha-1(tm329)* mutant animals. We observed a moderate reduction in fertility in *rha-1(tm329)* mutants at normal growth temperature (20 °C) (fig. S1A). When grown at 25 °C, *rha-1(tm329)* mutants exhibited a progressive decline in fertility and were sterile by the second generation (Fig. 1C). These results suggest RHA-1 promotes fertility and germline immortality, consistent with a role in germline small RNA pathways.

Next, we investigated whether RHA-1 ATPase activity contributes to RHA-1 function in RNAi and fertility. To this aim, we generated two RHA-1 ATPase activity mutants by substituting critical amino acids in the conserved ATP binding and ATP hydrolysis domains of DExH family helicases (*32*, *33*). In the background of endogenously tagged GFP::RHA-1, we used CRISPR Cas9-mediated gene editing to substitute the critical lysine residue in the GKT motif required for ATP binding with glutamine, generating an ATP binding mutant *gfp::rha-1(K410Q)*. Additionally, we substituted the key glutamic acid residue of the ATP hydrolysis domain with alanine, generating an ATP hydrolysis mutant *gfp::rha-1(E505A)*. First, we examined whether RHA-1 ATPase activity regulates RNAi as observed for *rha-1(tm329)* mutants. In both RHA-1 ATP binding and ATP hydrolysis mutants, RNAi of germline gene *pos-1* or *mex-3* was much less effective in comparison to control animals (Fig. 1D). Like *rha-1(tm329)* mutants, RHA-1 ATP binding and ATP hydrolysis mutants were fully sensitive to RNAi of somatic genes *dpy-13* and *unc-22* (Fig. 1E). These data suggest RHA-1 ATPase activity contributes to RHA-1 function in germline RNAi.

Next, we investigated whether RHA-1 ATPase activity regulates germline fertility, and immortality. At 20 °C, RHA-1 ATP binding and ATP hydrolysis mutants showed a modest reduction in fertility (fig. S1B). At 25 °C, RHA-1 ATP binding and ATP hydrolysis mutants exhibited progressively reduced fertility and reached near sterility by the second generation (Fig. 1F). Thus, loss of RHA-1 ATPase activity impacts germline fertility and immortality comparable to the complete loss of RHA-1, indicating RHA-1 ATPase activity is critical for these processes.

### RHA-1 localizes to germ cell nuclei and germ granules

Based on our findings that RHA-1 performs a role in germline RNAi and fertility, we investigated RHA-1 localization. To this aim, we introduced a GFP tag at the N-terminus of endogenous RHA-1 using CRISPR Cas9-mediated gene editing. We find RHA-1 is expressed in somatic and germ cells throughout development, including the spermatogenic germline of L4 hermaphrodites and males, the oogenic germline of adult hermaphrodites, and developing embryos (Fig. 2A and fig. S2, A and B). In germ cells, comparing RHA-1 localization to nuclear pore protein NPP-7, we find RHA-1 localizes both inside and outside of nuclei in puncta that resemble germ granules (Fig. 2A).

**Figure 2.**
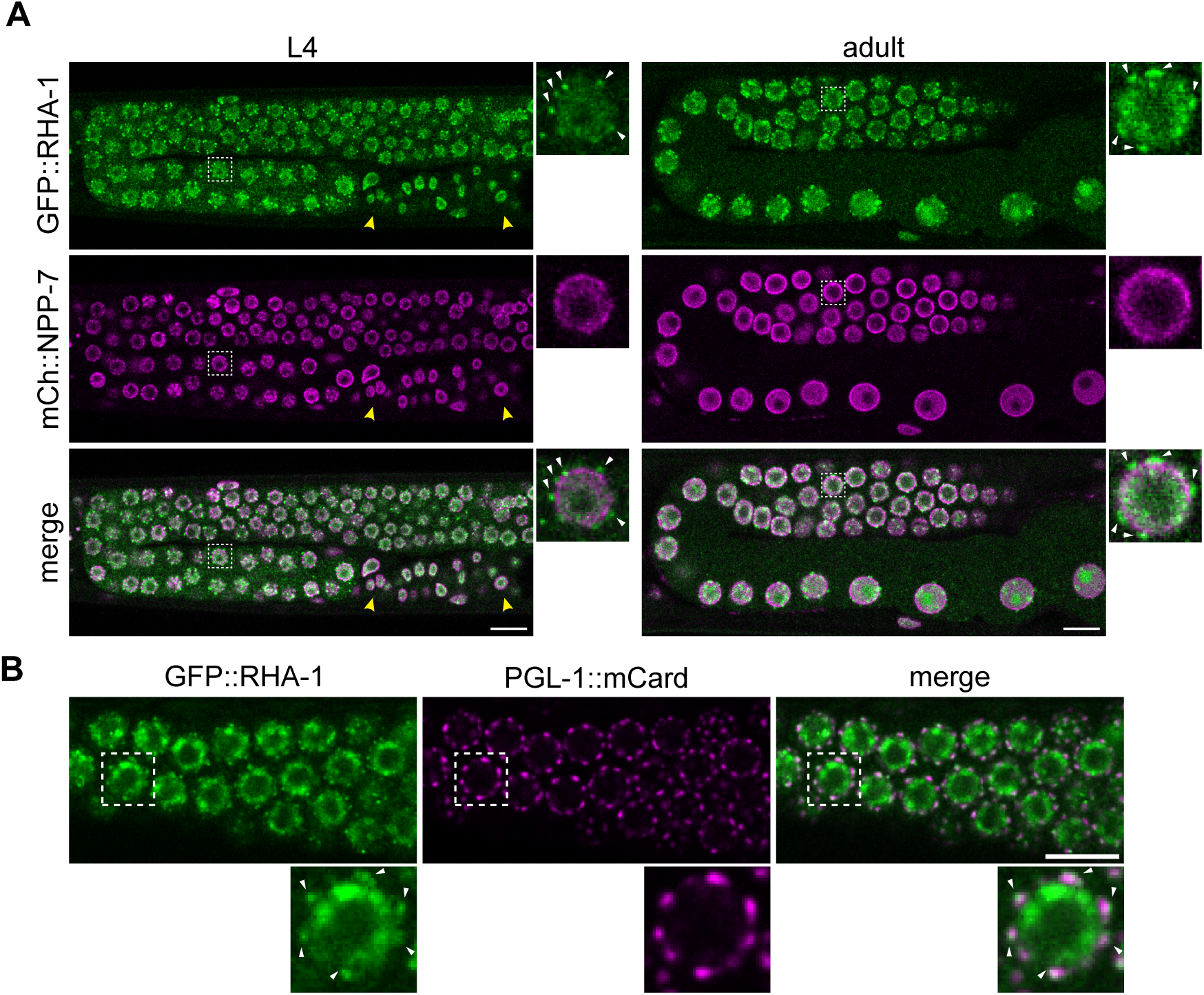
RHA-1 localizes to germ cell nuclei and germ granules. (**A**) Representative live-fluorescent images of animals expressing GFP::RHA-1 and mCherry:NPP-7 (nuclear pore marker). L4 germline (left) and adult germline (right). Boxes with dashed outline show the single cropped and enlarged nucleus. Between the yellow arrowheads are somatic cell nuclei outside of the germline. (**B**) Representative live-fluorescent images of adult pachytene region germ cells in animals expressing GFP::RHA-1 and PGL-1::mCardinal (P compartment marker). (**A** and **B**) Boxes with dashed outline show the single cropped and enlarged nucleus. White arrowheads point to the RHA-1 granules at the nuclear periphery outside of the germ cell nucleus. Scale bars = 10 µm.

Germ granules are RNA-rich biomolecular condensates that contain small RNA factors and RNA processing machinery (*34–36*).To determine whether RHA-1 localizes to germ granules, we examined RHA-1 localization relative to PGL-1-marked germ granules. We find RHA-1 granules are in proximity but not fully overlapped with PGL-1-marked granules surrounding germ cell nuclei (Fig. 2B). This result suggests RHA-1 and PGL-1 localize to distinct sub-compartments of germ granules (see additional details below). Nonetheless, our observation identifies RHA-1 as a germ granule factor throughout germ cell development.

### RHA-1 promotes 22G-RNA production and sorting from CSR-1 target genes

Given several germ cell-only RNAi factors, including EGO-1 and CSR-1, contribute to 22G-RNA production, this raised the possibility that RHA-1 is also involved in 22G-RNA production. We performed deep sequencing of the small RNA species isolated from young adult wild type and *rha-1(tm329)* mutants grown at normal growth temperature (20 °C). We first assessed the overall abundance of the major classes of small RNAs including piRNAs, miRNAs, and 22G-RNAs. In result, neither piRNAs nor miRNAs showed a notable change in abundance in *rha-1(tm329)* mutants compared to wild type across replicates (fig. S3, A and B). Additionally, the abundance of 22G-RNAs mapping to WAGO and CSR-1 target genes showed no uniform change in *rha-1(tm329)* mutants compared to wild type (Fig. 3A and fig. S3C).

**Figure 3.**
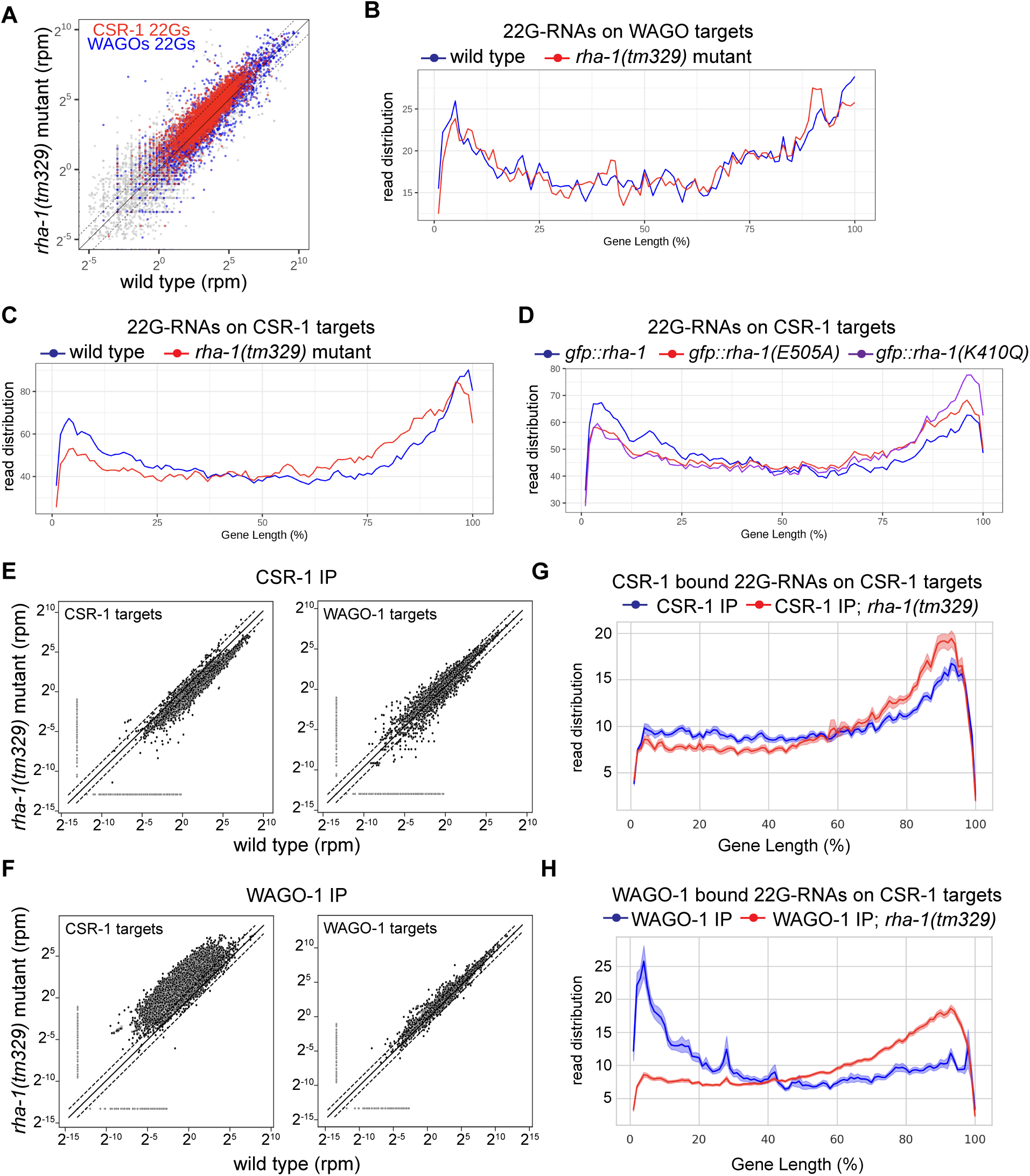
RHA-1 is required for the biogenesis of CSR-1 bound 22G-RNAs and 22G-RNA sorting. (**A**) Scatterplot showing the abundance of all 22G-RNAs mapped to each CSR-1 target (red) and WAGO target (blue) in wild type worms compared to *rha-1(tm329)* mutant worms. The three diagonal lines indicate a two-fold increase (top), no change (middle), or a two-fold depletion (bottom) in *rha-1(tm329)* mutant. (**B** and **C**) Metagene traces show the distribution of normalized 22G-RNAs (sRNA-seq) mapping to WAGO (**B**) or CSR-1 (**C**) target genes by percentage of WAGO or CSR-1 target gene length. Reads from wild type (blue) and *rha-1(tm329)* mutant (red). (**D**) Metagene traces show the distribution of normalized 22G-RNAs (sRNA-seq) mapping to CSR-1 target genes by percentage of CSR-1 target gene length. Reads from control *gfp::rha-1* (blue), *gfp::rha-1(E505A)* (red), and *gfp::rha-1(K410Q)* (purple) animals. (**E** and **F**) Scatterplots showing the abundance of CSR-1 bound 22G-RNAs (CSR-1 IP-sRNA-seq) (**E**) or WAGO-1 bound 22G-RNAs (WAGO-1 IP-sRNA-seq) (**F**) mapped to each CSR-1 target (left) and WAGO target (right) in wild type worms compared to *rha-1(tm329)* mutant worms. The three diagonal lines indicate a two-fold increase (top), no change (middle), or a two-fold depletion (bottom) in *rha-1(tm329)* mutant. (**G** and **H**) Metagene traces show the read counts distribution of normalized CSR-1 bound 22G-RNAs (IP-sRNA-seq) (**G**) or WAGO-1 bound 22G-RNAs (WAGO-1 IP-sRNA-seq) (**H**) mapping to CSR-1 target genes by percentage of CSR-1 target gene length. Traces show the mean read counts +/- one standard error. Reads from wild type IPs (blue) and *rha-1(tm329)* mutant IP samples (red).

22G-RNAs are produced by RdRPs directly from mRNA targets and processive RNA helicases such as DRH-3 and ZNFX-1 facilitate the spread of 22G-RNA production across the entire length of the mRNA targets (*37*, *38*). Given RHA-1 is similarly a processive RNA helicase, we analyzed the 22G-RNA distribution on WAGO and CSR-1 target genes. For WAGO target genes, the 22G-RNA distributions were similar for *rha-1(tm329)* mutants and wild type (Fig. 3B and fig. S3D). Strikingly, the 22G-RNA distributions on CSR-1 target genes showed an increase in production from the 3’ end with a drop off in production at the 5’ end of gene lengths in *rha-1(tm329)* mutants compared to wild type (Fig. 3C and fig. S3E). These findings suggest RHA-1 contributes to the production of 22G-RNAs derived across the full length of CSR-1 target mRNAs. Consistent with a role for helicase activity in this 22G-RNA production, we find RHA-1 ATP binding and ATP hydrolysis mutants show a similar change to the 22G-RNA distribution on CSR-1 target genes as *rha-1(tm329)* mutant animals (Fig. 3D). These data indicate RHA-1 helicase activity facilitates 22G-RNA production across the full length of CSR-1 target mRNAs.

Since 22G-RNAs are bound by CSR-1 and WAGOs, we asked whether the changes in 22G-RNA distribution on CSR-1 target genes in *rha-1(tm329)* mutants reflect changes to CSR-1 bound 22G-RNAs. To examine this, we immunoprecipitated either CSR-1 or WAGO-1 and sequenced the bound small RNAs with the expected outcome that CSR-1 22G-RNAs would report the same changes observed in total small RNAs, while WAGO-1 22G-RNAs would report no change. Instead, we found changes in the abundance of both CSR-1 and WAGO-1 bound 22G-RNAs derived from CSR-1 target genes. For CSR-1 bound 22G-RNAs, we find reduced overall abundance mapping to CSR-1 target genes (Fig 3E). Strikingly, for WAGO-1 bound 22G-RNAs, we find increased abundance mapping to CSR-1 target genes (Fig. 3F). These observations suggest that RHA-1 promotes CSR-1 22G-RNA biogenesis and prevents the aberrant binding of CSR-1 target derived 22G-RNAs by WAGO-1.

Although WAGO-1 aberrantly binds small RNAs mapping to CSR-1 target genes in *rha-1(tm329)* mutants, WAGO-1 bound 22G-RNAs mapping to WAGO target genes showed no change in abundance from wild type (Fig. 3F). Additionally, CSR-1 bound 22G-RNAs mapping to WAGO target genes showed no change in abundance for *rha-1(tm329)* mutants compared to wild type (Fig. 3E). These results suggest that RHA-1 does not regulate the biogenesis or sorting of 22G-RNAs derived from WAGO target genes.

The distributions of CSR-1 and WAGO-1 bound 22G-RNAs on CSR-1 target genes showed decreased abundance of small RNAs derived from the 5’ end of mRNAs in *rha-1(tm329)* mutants compared to wild type, consistent with our results from total small RNAs (Fig. 3, G and H). Inspection of the CSR-1 bound 22G-RNAs mapping to several individual CSR-1 target genes showed CSR-1 22G-RNAs greatly reduced from the 5’ region of genes in *rha-1(tm329)* mutants compared to wild type (fig. S4A). For these CSR-1 target genes, the incorrectly sorted WAGO-1 bound 22G-RNAs in *rha-1(tm329)* mutants are also reduced at the 5’ end of genes (fig. S4A). In contrast, WAGO-1 and CSR-1 bound 22G-RNAs mapping to several WAGO target genes showed relatively no change in *rha-1(tm329)* mutants compared to wild type (fig. S4B). This further supports our findings that RHA-1 regulates 22G-RNA biogenesis and sorting from CSR-1 target mRNAs, but not WAGO target mRNAs.

We next aimed to gain insight into how RHA-1 contributes to CSR-1 22G-RNA distribution. Notably, both *rha-1(tm329)* and CSR-1 slicer mutants show similar changes to the CSR-1 bound 22G-RNA distribution on CSR-1 target gene lengths, where 22G-RNA production from the 5’ end of mRNAs is greatly reduced (fig. S5, A and B). Given that CSR-1-mediated cleavage of target mRNAs is proposed to initiate 22G-RNA synthesis from the new cleaved 3′ ends, sequential CSR-1 slicing events are thought to drive the production of 22G-RNAs by EGO-1 from the 3’ end toward the 5′ end across the full length of target mRNAs (*23*). Like *rha-1* mutants, CSR-1 slicer mutants showed no change in total abundance of 22G-RNAs mapped to CSR-1 targets but showed a slight decrease in abundance of CSR-1 bound 22G-RNAs mapped to CSR-1 targets (fig. S5, C and D). This hints that small RNAs may also be mis-sorted in CSR-1 slicer mutants. Together, these data suggest both RHA-1 and CSR-1 slicer activity are critical for 22G-RNA production from the 5’ region of target mRNAs. Our analyses raise the possibility that RHA-1 may promote CSR-1 slicer activity or facilitate sequential rounds of CSR-1-mediated cleavage and 22G-RNA synthesis on target mRNAs.

Overall, these data lead us to propose that RHA-1 promotes 22G-RNA biogenesis from the 5’ region of CSR-1 target mRNAs and 22G-RNA sorting to CSR-1.

### RHA-1 localizes to the E compartment of germ granules

Germ granules exhibit a complex organization of spatially distinct sub-compartments (*39–42*). These sub-compartments are enriched for distinct sets of small RNA components and generally linked to specific pathway functions. Specifically, the RdRP EGO-1, which is responsible for CSR-1 22G-RNA biogenesis, is largely enriched in the E compartment, while CSR-1 and WAGO-1 are enriched in D and P compartments, respectively (Fig. 4A) (*43*). Based on our results that RHA-1 promotes CSR-1 22G-RNA biogenesis, we wondered whether RHA-1 localizes within the E compartment of germ granules. Thus, we examined RHA-1 localization compared to EGO-1 and found RHA-1 granules highly overlap with EGO-1 (Fig. 4B). In addition, we observed that RHA-1 exhibited higher levels of co-localization with E compartment factor ELLI-1, than with P compartment factor PGL-1 (Fig. 4, C and D).

**Figure 4.**
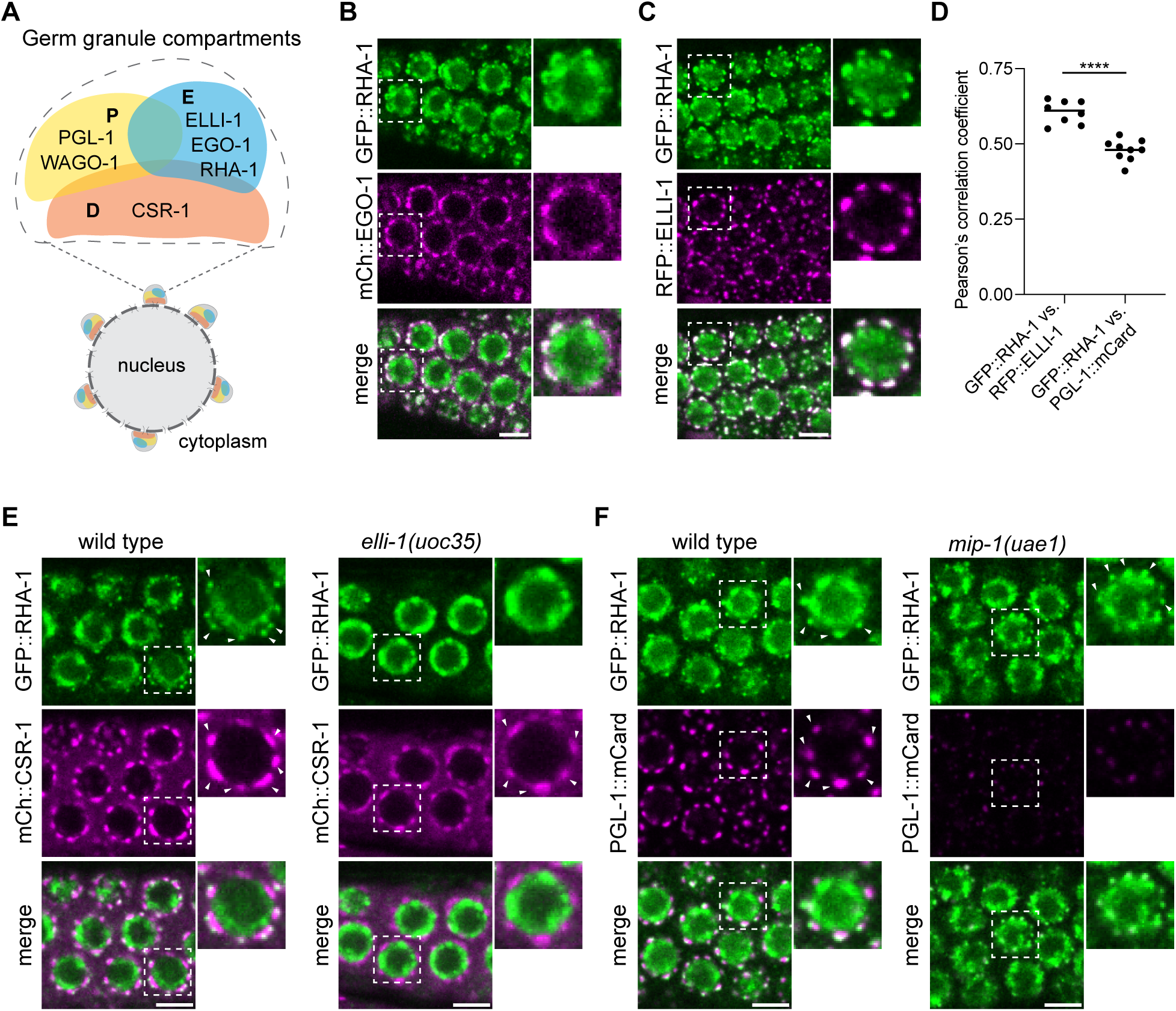
RHA-1 localizes to the E compartment of germ granules. (**A**) Cartoon depicting germ granule sub-compartment spatial organization and factors that localize to these sub-compartments. (**B** and **C**) Representative live-fluorescent images of adult germ cells in the pachytene region in animals expressing GFP::RHA-1 and mCherry::EGO-1 (E compartment marker) (**B**) or RFP::ELLI-1 (E compartment marker) (**C**). (**D**) Quantification of colocalization between the indicated fluorescent proteins of pachytene germ cells. Each data point represents the Pearson’s R-value showing the degree of colocalization between two fluorescence channels indicated. The solid black line indicates the mean value. Each data point represents independent gonad images. *N =* 8-9 gonads. Two-tailed p values were calculated using Mann-Whitney-Wilcoxon test. ****, p<0.0001. (**E**) Representative live-fluorescent images of pachytene germ cells show the localization of GFP::RHA-1 and mCherry::CSR-1 (D compartment marker) in wild type (left) or *elli-1(uoc35)* mutant animals (right). White arrowheads point to the RHA-1 or CSR-1 granules at the nuclear periphery outside of the germ cell nucleus. (**F**) Representative live-fluorescent images of pachytene germ cells show the localization of GFP::RHA-1 and PGL-1::mCardinal (P compartment marker) in wild type (left) or *mip-1(uae1)* mutant animals (right). White arrowheads point to the RHA-1 or PGL-1 granules at the nuclear periphery outside of the germ cell nucleus. For all images, boxes with dashed outline show the single cropped and enlarged nucleus. Scale bars = 5 µm.

The E compartment of germ granules specifically requires the assembly factor ELLI-1 for formation (*44*). To examine whether RHA-1 requires the E compartment for localization to germ granules, we examined RHA-1 localization in *elli-1(uoc35)* mutants. Strikingly, in *elli-1(uoc35)* mutants, RHA-1 germ granule localization is completely disrupted, while D compartment factor CSR-1 germ granule localization is maintained (Fig. 4E). In contrast, loss of the P compartment assembly factor MIP-1 strongly disrupted P compartment factor PGL-1 germ granule localization but not RHA-1 granule localization (Fig. 4F). Together, these findings identify RHA-1 as an E compartment factor within germ granules (Fig. 4A). Intriguingly, loss of E compartment assembly was recently shown to disrupt EGO-1 22G-RNA production specifically from the 5’ region of mRNAs (*44*). Together, these results suggest that RHA-1 and EGO-1 enriched within the E compartment facilitate 22G-RNA production from the 5’ region of CSR-1 target mRNAs.

### RHA-1 localization to the E compartment depends on CSR-1 and EGO-1

The organization of small RNA factors within germ granule compartments is proposed to facilitate the fidelity of gene regulation by small RNAs (*42*, *45*, *46*). Given the observed changes in 22G-RNA biogenesis and sorting in *rha-1* mutants, we wondered whether loss of RHA-1 leads to disrupted germ granule compartment formation or organization. We focused on the P, D, and E sub-compartments of germ granules since WAGO-1, CSR-1, and EGO-1 localize to these compartments respectively. Fluorescence imaging of PGL-1-marked P compartments and EGO-1- and ELLI-1-marked E compartments revealed a slight increase to granule size in *rha-1(tm329)* mutants compared to wild type (Fig. 5, A to D). For WAGO-1-marked P compartments and CSR-1-marked D compartments, we observed no change to granule size (Fig. 5, A and B). These results suggest RHA-1 does not greatly impact the formation of these compartments.

**Figure 5.**
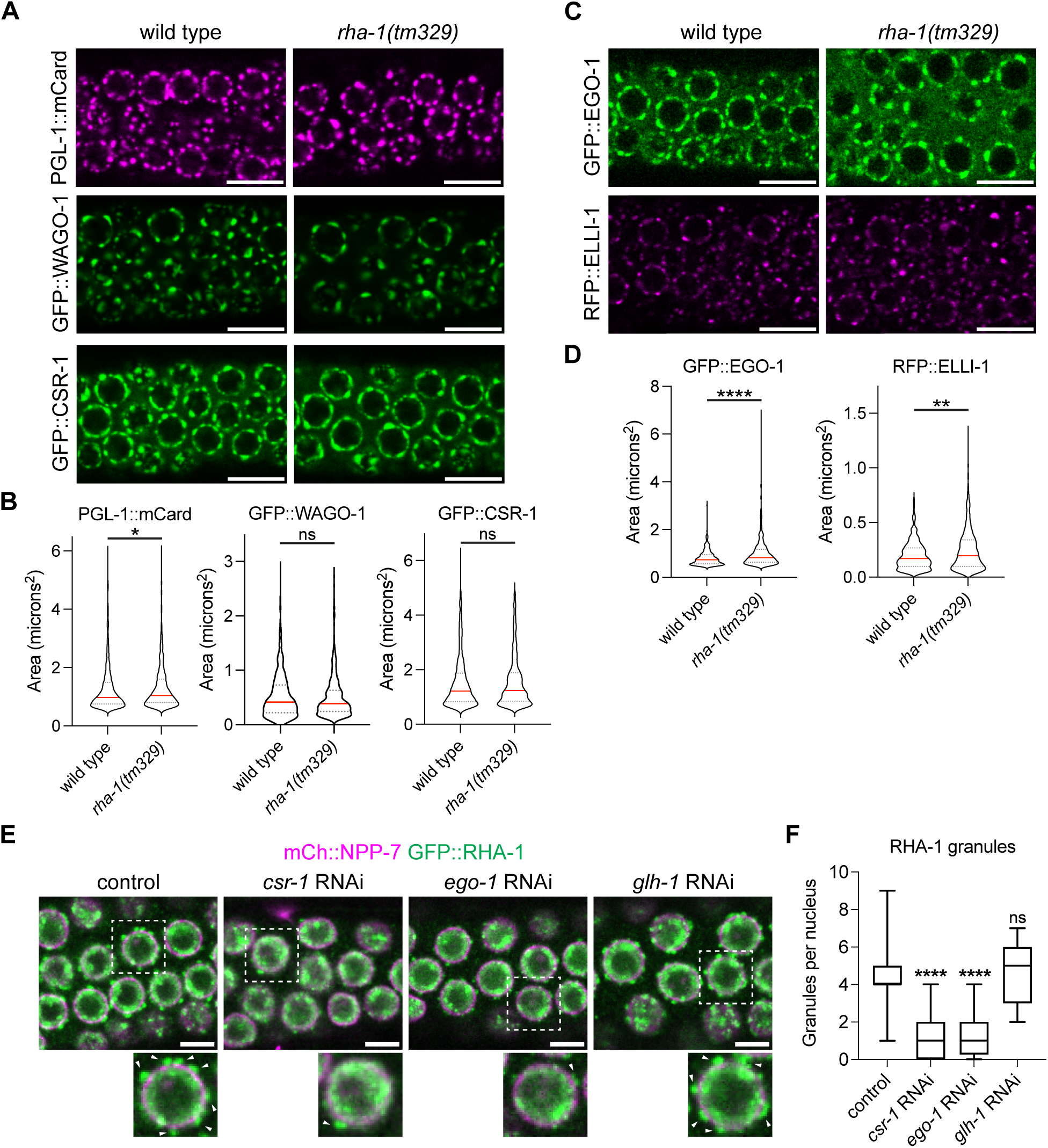
RHA-1 germ granule localization depends on CSR-1 and EGO-1. (**A** and **C**) Representative live-fluorescent images of pachytene germ cells show the localization of the indicated germ granule markers in wild type and *rha-1(tm329)* mutant animals. (**A**) PGL-1::mCard and GFP::WAGO-1 are P compartment markers, and GFP::CSR-1 is the D compartment marker. (**C**) GFP::EGO-1 and RFP::ELLI-1 are E compartment markers. Scale bars = 10 µm. (**B** and **D**) Violin plots depicting granule size (µm^2^) quantified from experiment in part (**A** and **C**) for *N* = 8-10 independent germlines for wild type and *rha-1(tm329)* mutant animals. Red line marks the median value. Two-tailed p values were calculated using Mann-Whitney-Wilcoxon test. *, p<0.05; **, p<0.01; ***, p<0.001. ns, no significance. (**E**) Representative live-fluorescent merged images of pachytene germ cells in animals expressing GFP::RHA-1 (green) and mCherry:NPP-7 (nuclear pore marker) (magenta) in control or RNAi knockdown of the indicated genes. Boxes with dashed outline show the single cropped and enlarged nucleus. White arrowheads point to RHA-1 granules. Scale bars = 5 µm. (**F**) Boxplots show the number of GFP::RHA-1 granules per germ cell nuclei from the experiment in (**E**). *N =* 8-10 gonads and number of granules per nucleus was quantified for 10 nuclei per gonad. For the boxplots, the line indicates the median value, the box indicates the first and third quartiles, and the whiskers indicate the min and max values. Two-tailed p values were calculated using Mann-Whitney-Wilcoxon test. ****, p<0.001. ns, no significance.

Next, we examined whether loss of RHA-1 leads to changes in germ granule compartment organization. Typically, the D compartment localizes closest to the nuclear envelope with the P and E compartments positioned adjacent to each other just outside of the D compartment (Fig. 4A) (*40*). These compartments also partially overlap indicative of factor interaction between sub-compartments. We compared the relative positions of these compartments by pair-wise comparisons in co-marked strains. Overall, the relative positions of the D, P, and E compartments showed similar organization in *rha-1(tm329)* mutants as wild type (fig. S6, A to C). We measured a decrease in co-localization between the P and D compartments and P and E compartments in *rha-1(tm329)* mutants compared to wild type (fig. S6, D and E). Co-localization between D and E compartments showed no change (fig. S6F). We also noted that the increase in granule sizes for PGL-1 and EGO-1 in *rha-1(tm329)* mutants were not always apparent in these co-marked backgrounds (fig. S6, A to C). This data suggests loss of RHA-1 leads to minor changes in germ granule compartment organization.

Following from our findings that RHA-1 does not greatly disrupt germ granule formation or organization, we hypothesized RHA-1 may function in CSR-1 22G-RNA biogenesis downstream of CSR-1 or EGO-1 activity. To test whether RHA-1 localization depends on CSR-1 or EGO-1, we performed RNAi knockdown of *csr-1* or *ego-1*, then evaluated RHA-1 localization compared to nuclear pore marker NPP-7. As a result of both *csr-1* or *ego-1* knockdown, RHA-1 germ granule localization was largely disrupted and formed a reduced number of germ granules per nucleus (Fig. 5, E and F). In contrast, knockdown of *glh-1*, a factor well known for its role in P granule assembly, resulted in no change to RHA-1 germ granule localization or granule number per nucleus (Fig. 5, E and F) (*46*, *47*). These results indicate RHA-1 localization to germ granules depends on CSR-1 and EGO-1, but not P granule factor GLH-1. Moreover, these results indicate RHA-1 germ granule recruitment downstream of CSR-1 or EGO-1 may contribute to RHA-1 specificity in small RNA biogenesis on CSR-1 target mRNAs.

### CSR-1 target genes are upregulated in *rha-1* mutants

Since small RNAs regulate target gene expression, we next examined whether *rha-1(tm329)* mutants exhibit changes in gene expression, particularly for CSR-1 target genes. To this aim, we generated mRNA high-throughput sequencing libraries from young adult wild type and *rha-1(tm329)* mutants grown at 20 °C and evaluated mRNA level fold changes. We proposed two potential outcomes based on our small RNA results. First, incorrect sorting of 22G-RNAs to WAGO-1 could lead to WAGO-1-mediated RNA silencing, and as a result, CSR-1 target mRNA levels would be reduced in *rha-1(tm329)* mutants compared to wild type. Alternatively, reduction in CSR-1 bound 22G-RNAs could lead to loss of CSR-1-mediated mRNA fine-tuning, and as a result, CSR-1 target mRNA levels would be increased in *rha-1(tm329)* mutants. The mRNA sequencing results displayed an increase in mRNA fold change for CSR-1 targets in *rha-1(tm329)* mutants. In contrast, WAGO target genes showed relatively no change (Fig. 6A). Additionally, we evaluated CSR-1 top slicer target genes defined in a previous report as genes with 2-fold or more decrease in CSR-1 IP-associated 22G-RNAs in the CSR-1 slicer mutant, and 2-fold or more increase in mRNA level in the CSR-1 slicer mutant compared to wild type (*23*). The CSR-1 top slicer targets showed an increase in mRNA fold change in *rha-1(tm329)* mutants compared to wild type at a higher fold than all CSR-1 targets (Fig. 6A). This data suggests that loss of RHA-1 mainly disrupts CSR-1-mediated mRNA fine-tuning. Additionally, in *rha-1(tm329)* mutants the gained WAGO-1 bound 22G-RNAs mapping to CSR-1 target genes do not aberrantly silence these targets.

**Figure 6.**
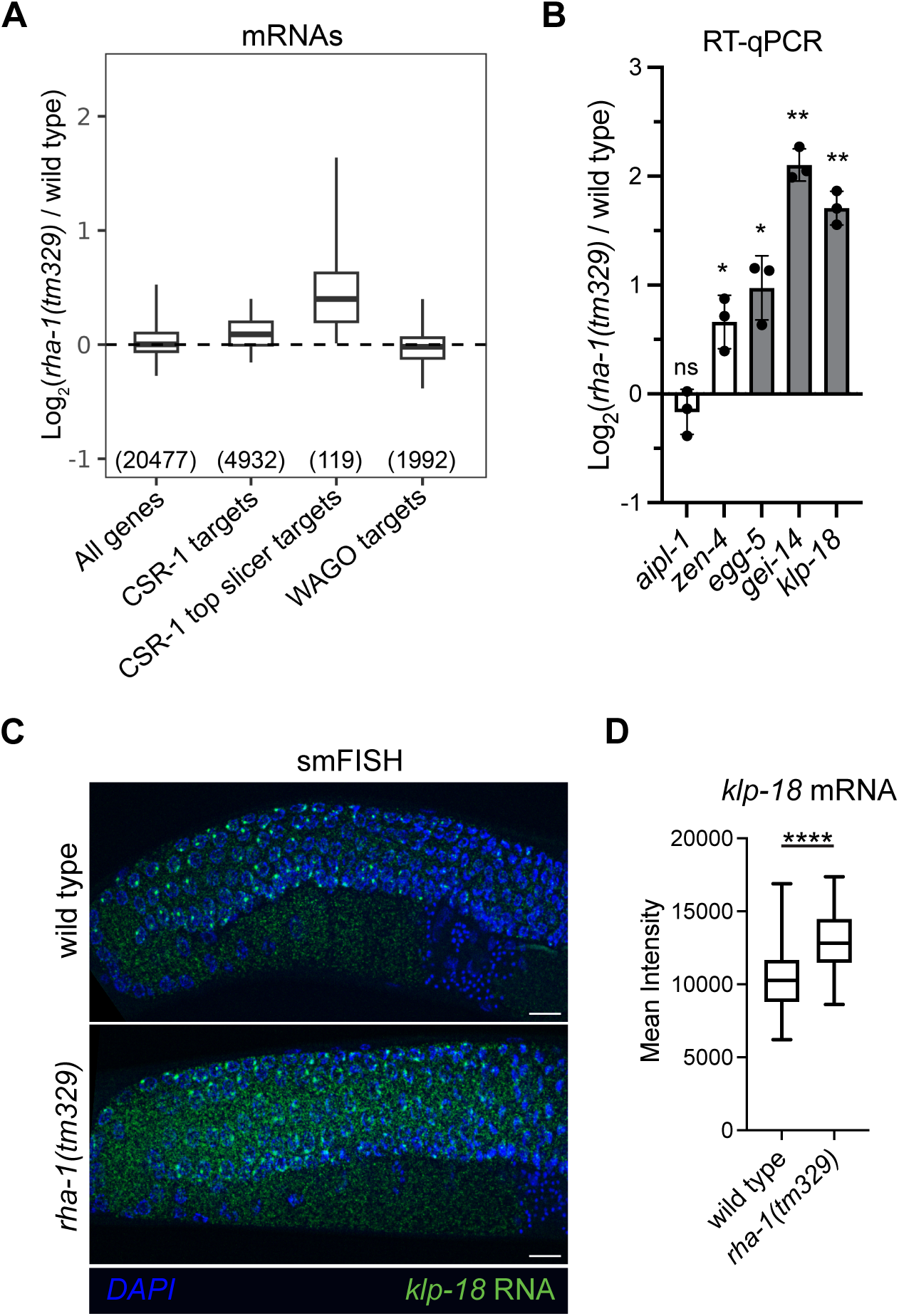
CSR-1 target mRNAs are upregulated in *rha-1* mutants. (**A**) Boxplot showing log_2_ fold change in mRNA abundance for adult stage *rha-1(tm329)* mutants compared to wild type animals (two biological replicates). For the boxplot, the line indicates the median value, the box indicates the first and third quartiles, and the whiskers indicate the 5th and 95th percentiles, excluding outliers. The sample size n (genes) is indicated in parentheses. (**B**) Bar graph showing log_2_ fold change in mRNA expression (RT-qPCR) of the indicated genes in wild type compared to *rha-1(tm329)* mutant young adult animals. White bars are CSR-1 targets and gray bars indicate CSR-1 top slicer targets. Data are represented as mean and standard deviation for three biological replicates. Two-tailed p values were calculated from ΔCt values using Mann-Whitney-Wilcoxon test. *, p<0.05; **, p<0.01. ns, no significance. (**C**) Representative images of *klp-18* mRNA smFISH in wild type and *rha-1(tm329)* mutant adult germlines. DAPI stain (blue) and *klp-18* mRNA (green). Scale bars = 10 µm. (**D**) Boxplot depicting smFISH quantification of *klp-18* mRNA in wild type and *rha-1(tm329)* mutant adult germlines. *N =* 10-12 independent animals. For the boxplot, the line indicates the median value, the box indicates the first and third quartiles, and the whiskers indicate the minimum and maximum. Two-tailed p values were calculated using Mann-Whitney-Wilcoxon test. ****, p<0.0001.

Given loss of RHA-1 largely impacts CSR-1 bound small RNAs and regulation of target genes, we compared the genes that exhibit mRNA level changes in *rha-1(tm329)* mutants to those in CSR-1 compromised mutants. We performed DESeq2 analysis to identify significantly upregulated and downregulated mRNAs in *rha-1(tm329)* mutants, CSR-1 slicer mutants, and CSR-1 depleted animals (*21*, *48*). Of the genes that exhibit significantly upregulated mRNA levels in *rha-1(tm329)* mutants (DESeq2 adjusted p value of ≤0.05), 63.1% are shared with those in CSR-1 slicer mutants, CSR-1 depleted animals, or both (fig. S7A). In contrast, none of the significantly downregulated mRNAs in *rha-1(tm329)* mutants are in common with either CSR-1 slicer mutants or CSR-1 depleted animals (fig. S7B). Taken together, this data suggests RHA-1 activity is important for fine-tuning the mRNA level of CSR-1 target genes, like CSR-1 slicer activity.

To validate our mRNA sequencing results, we evaluated mRNA level fold change by RT-qPCR for several CSR-1 and CSR-1 top slicer target genes. CSR-1 target genes *aipl-1* and *zen-4* showed a slight decrease or increase in mRNA expression respectively (Fig. 6B). CSR-1 top slicer target genes *egg-5*, *gei-14*, and *klp-18*, all showed a nearly 2-fold or more increase mRNA expression (Fig. 6B). Additionally, we performed single molecule fluorescence in situ hybridization (smFISH) for CSR-1 top slicer target *klp-18* mRNA in young adults. We observed an increase in *klp-18* mRNA signal throughout the adult syncytial germline in *rha-1(tm329)* mutants compared to wild type (Fig. 6, C and D). We also performed smFISH for *klp-18* in CSR-1 depleted animals utilizing a CSR-1 degron strain (*22*). Like *rha-1(tm329)* mutants, CSR-1 depleted animals displayed an increase in *klp-18* mRNA signal throughout the adult germline compared to control animals (fig. S7, C and D).

Overall, we find loss of RHA-1 leads to increased mRNA levels of CSR-1 target genes, especially CSR-1 slicer target genes, suggesting RHA-1 promotes CSR-1 target mRNA fine-tuning.

### RHA-1 safeguards male fertility and sperm differentiation

CSR-1 small RNAs target germline-expressed genes throughout development, including mRNAs enriched during spermatogenesis and oogenesis(*29*, *49*). At the earlier L4 stage, germ cells actively undergo spermatogenesis, and by the adult stage, germ cells switch to oocyte production. Interestingly, for young adult animals grown at 20 °C we found increased mRNA levels for spermatogenesis-enriched genes in *rha-1(tm329)* mutants compared to wild type, while oogenesis-enriched genes show relatively no change (Fig. 7A). Since spermatogenic genes are no longer transcribed in adults, the increase in spermatogenic mRNAs in *rha-1(tm329)* mutant animals likely results from a defect in post-transcriptional mRNA degradation. In follow up, we performed smFISH for spermatogenic gene *gipc-1* mRNA in L4 stage animals grown at 20 °C. We found increased levels of *gipc-1* mRNA in *rha-1(tm329)* mutant germlines compared to wild type (Fig. 7B). This data indicates RHA-1 promotes downregulation of spermatogenic mRNAs in germ cells actively undergoing spermatogenesis

Since *rha-1(tm329)* mutants showed elevated spermatogenic mRNA levels, we investigated whether RHA-1 contributes to male fertility, female fertility, or both. For this, we utilized *rha-1(tm329); fog-2(q71)* double mutant animals. This background generates a population of animals producing only female germ cells and males producing normal male germ cells. Therefore, all offspring of this background result from mating, not self-fertilization. At 20 °C, mating between *rha-1(tm329); fog-2(q71)* mutant males and females resulted in a significant decrease in cross progeny (Fig. 7C). At 25 °C, mating of either *rha-1(tm329); fog-2(q71)* mutant females or males to *fog-2(q71)* animals resulted in a reduced number of cross progeny (Fig. 7D). Moreover, mating of *rha-1(tm329); fog-2(q71)* mutant females to *rha-1(tm329); fog-2(q71)* mutant males resulted in the smallest brood sizes (Fig. 7D). These mating results suggest RHA-1 promotes the fertility of both female and male germ cells.

**Figure 7.**
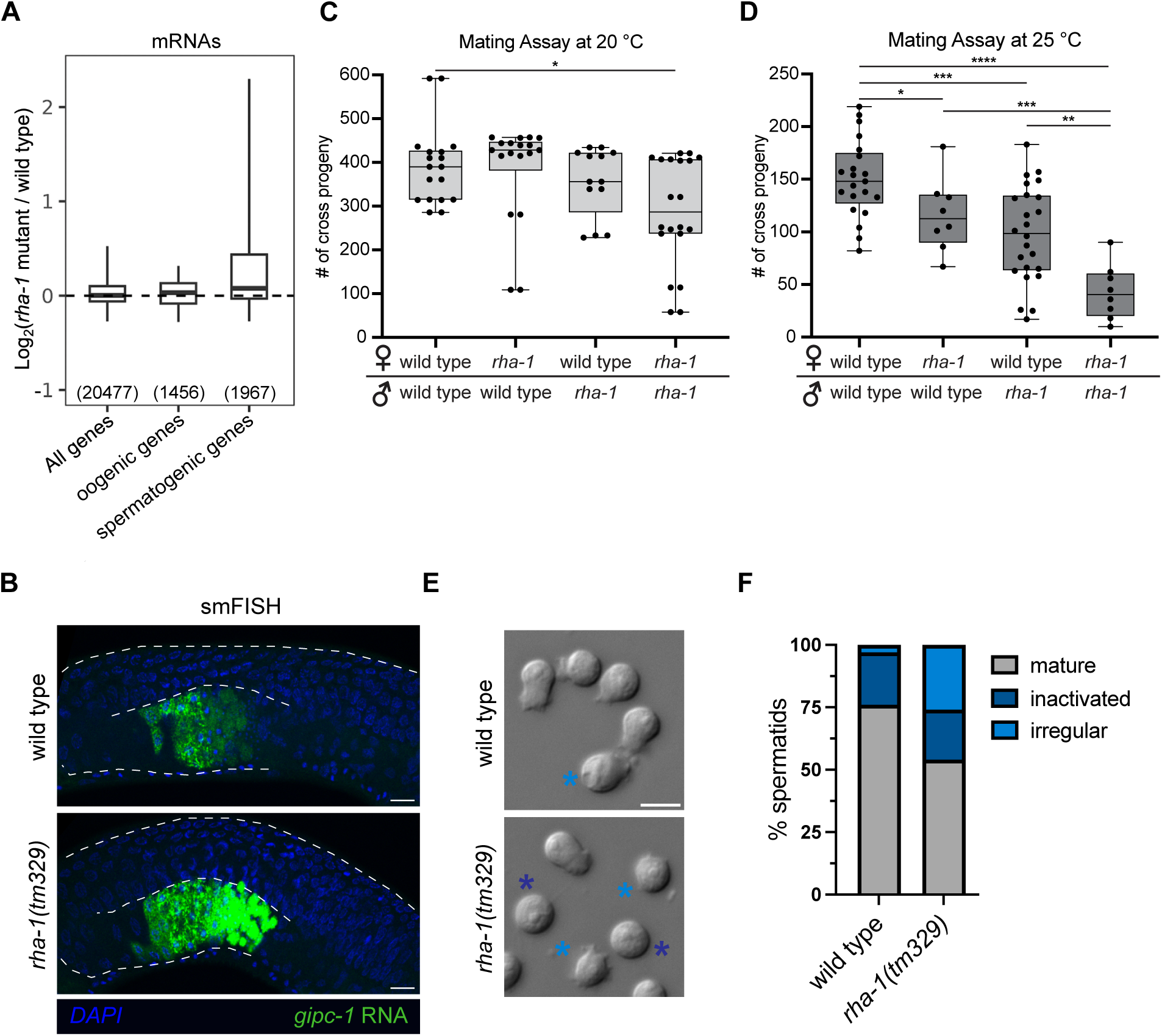
RHA-1 is required for optimal fertility and sperm differentiation. (**A**) Boxplot showing log_2_ fold change in mRNA abundance for adult stage *rha-1(tm329)* mutants compared to wild type animals (two biological replicates). For the boxplot, the line indicates the median value, the box indicates the first and third quartiles, and the whiskers indicate the 5th and 95th percentiles, excluding outliers. The sample size n (genes) is indicated in parentheses. (**B**) Representative images of *gipc-1* mRNA smFISH in L4 stage wild type and *rha-1(tm329)* mutant animals. DAPI stain (blue) and *gipc-1* mRNA (green). Dashed line marks the border of the germline. Scale bar = 10 µm. (**C** and **D**) Brood sizes from pair-wise mating between *fog-2(q71)* (wild type) and *rha-1(tm329); fog-2(q71)* mutant (*rha-1*) obligate females and males at (**C**) 20 °C or (**D**) 25°C. Data points correspond to the number of alive F1 cross progeny from individual obligate females. For the boxplots, the line indicates the median value, the box indicates the first and third quartiles, and the whiskers indicate the minimum and maximum. Two-tailed p values were calculated using Mann-Whitney-Wilcoxon test. *, p<0.05; **, p<0.01; ***, p<0.001; ****, p<0.0001. (**E**) Representative images of *in vitro* activated sperm by pronase treatment from wild type (top) and *rha-1(tm329)* mutant (bottom) animals. Dark blue asterisks mark inactivated round spermatids. Light blue asterisk mark irregular sperm. Scale bar = 5 µm. (**F**) Stacked bar graph showing percentage of activated mature, inactivated, and irregular spermatids from *in vitro* sperm activation assay for wild type and *rha-1(tm329)* mutant males where 10 adult male animals were dissected. Number of spermatids scored equals 833 for wild type and 857 for *rha-1(tm329)* mutant animals.

To investigate the cause of reduced male fertility observed in *rha-1* mutants, we examined sperm quality and function. During spermiogenesis, immature round spermatids differentiate into mature motile sperm with pseudopods. We isolated spermatids from *rha-1(tm329)* mutant and wild type males grown at 25 °C, and after inducing spermiogenesis *in vitro* by pronase treatment, we scored sperm for normal pseudopod formation, irregular pseudopod formation, or inactivated round spermatids (*50*). Most spermatids collected from wild type males developed normal pseudopods, with 24% combined irregular pseudopod formation or inactivated round spermatids (Fig. 7, E and F). Notably, just over half of spermatids collected from *rha-1(tm329)* mutant males developed normal pseudopods, with 46% combined irregular pseudopod formation or inactivated round spermatids (Fig. 7, E and F). Additionally, we found about half of the spermatids isolated from RHA-1 ATP binding and ATP hydrolysis mutant males were inactivated or formed irregular pseudopods, double that of the control male spermatids (fig. S8, A and B). Together, this data indicate RHA-1 is required for optimal male fertility and sperm differentiation.

Like RHA-1, CSR-1 has been shown to regulate male fertility and spermiogenesis (*51*). Additionally, RNAi-mediated knockdown of *csr-1* results in increased expression of spermatogenesis-enriched genes as measured by mRNA-sequencing (*52*). By smFISH, we also observed a strong increase in spermatogenic gene *gipc-1* mRNA level in L4 germlines of CSR-1 depleted compared to control animals (fig. S9A). We next wondered whether CSR-1 slicer activity contributes to CSR-1 function in male fertility. To test this, we utilized CSR-1 degron strains with transgenic rescue by either wild type slicer active CSR-1 or slicer dead CSR-1, or no transgenic rescue (*22*). Males for these three strains were grown without auxin for controls, or with auxin to deplete endogenous CSR-1 and mated to *fog-2(q71)* mutant obligate females at 20 °C (fig. S9B). Control males grown without auxin showed similar brood sizes for all three strains (fig. S9C). For males grown with auxin, brood sizes were significantly reduced for both CSR-1 slicer dead males and no rescue males compared to CSR-1 slicer active males (fig. S9C). This data indicates that like RHA-1, CSR-1 slicer activity also ensures optimal male fertility.

Overall, our data demonstrate loss of RHA-1 leads to elevated spermatogenic mRNA levels. This defect in mRNA regulation likely contributes to decreased male and female fertility observed in *rha-1* mutant animals. Additionally, our data hint at a critical gene regulatory role for RHA-1 ATPase and CSR-1 slicer activity during male germ cell development and differentiation.

## Discussion

A poorly understood aspect of small RNA networks is how each small RNA class establishes and regulates the correct mRNA targets. Here we identify a role for the conserved RNA helicase RHA-1 in regulating small RNA biogenesis and small RNA sorting to ensure proper gene expression in germ cells. More specifically, we find RHA-1 contributes to 22G-RNA production and sorting from CSR-1 target genes. Consistent with this role, RHA-1 localizes to E compartment of germ granules where factors that function in CSR-1 22G-RNA biogenesis are also located. Overall, the small RNA gene regulatory defects observed in *rha-1* mutants disrupt fertility, germline immortality, and sperm function, especially in elevated temperatures.

### Model for RHA-1 role in CSR-1 small RNA biogenesis and mRNA fine-tuning

An unresolved question is how CSR-1 performs opposing regulatory roles in mRNA fine-tuning and mRNA protection. One distinction between these two roles is that CSR-1 slicer activity mainly impacts CSR-1-mediated mRNA fine-tuning (*23*). CSR-1 slicer activity allows for amplified 22G-RNA production across the full length of target mRNAs but is dispensable for small RNA production at the 3’ ends. We find RHA-1 is similarly required for robust CSR-1 small RNA production from the 5’ regions of target mRNAs but dispensable for production at the 3’ ends (Fig. 8). Consistent with reduced abundance of CSR-1 22G-RNA targeting the full length of mRNAs, loss of RHA-1 leads to notable defects in mRNA fine-tuning.

**Figure 8.**
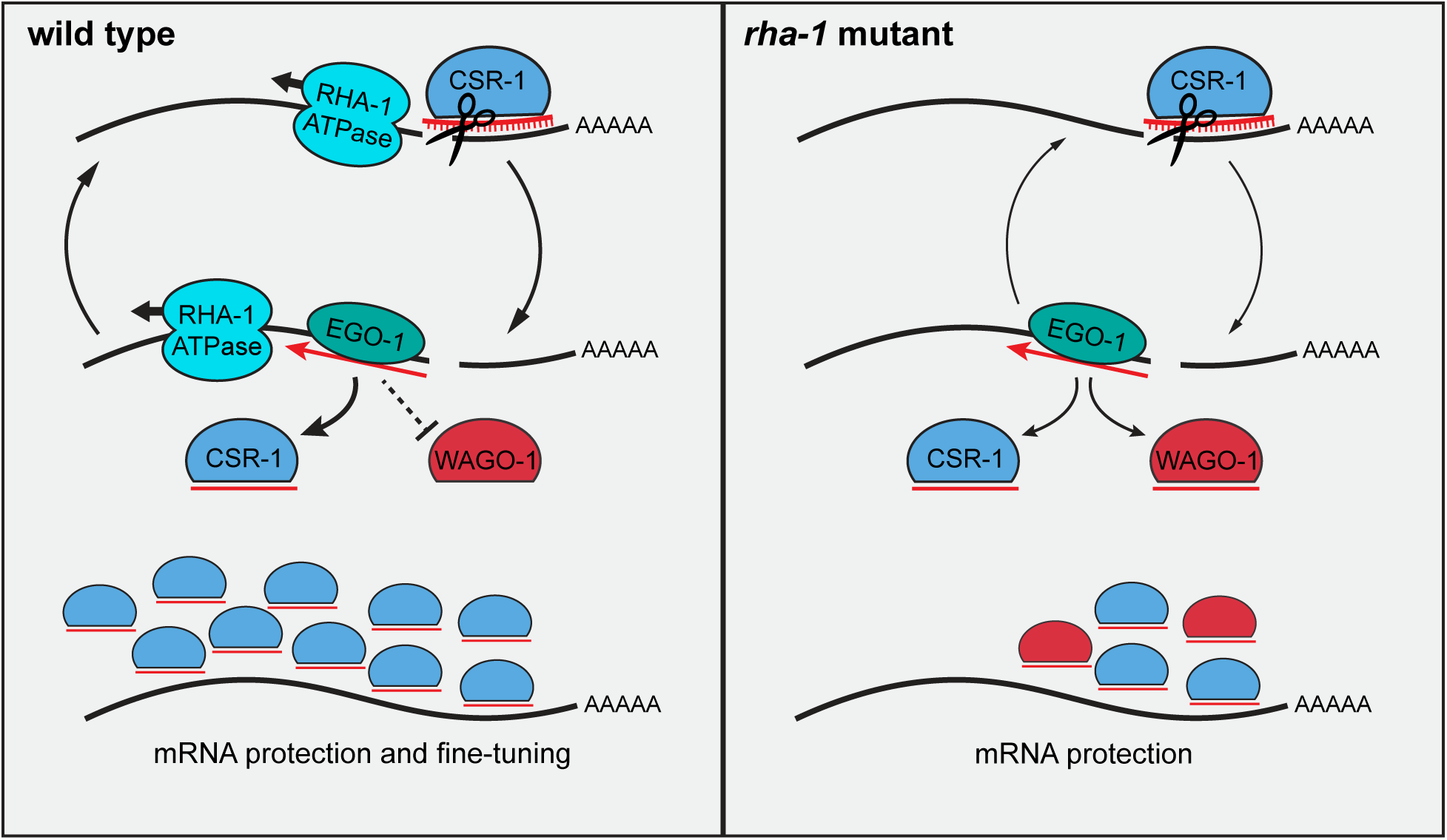
Model illustrating proposed function of RHA-1 in CSR-1 small RNA biogenesis and sorting. Model showing the proposed cycle of small RNA biogenesis on CSR-1 target mRNAs. (Left) RHA-1 ATPase activity may remodel RNP complexes on CSR-1 target mRNAs to facilitate multiple rounds of small RNA synthesis by EGO-1 and CSR-1 slicing. RHA-1 activity may promote the efficiency and mRNA availability during this cycle and facilitate small RNA sorting to CSR-1 over WAGO-1. CSR-1 targets are protected from WAGO-1 silencing and those targets with high abundance of CSR-1 bound small RNAs derived from the full mRNA length are fine-tuned by CSR-1. (Right) In *rha-1* mutants, the abundance of CSR-1 bound small RNAs is reduced from the 5’ region of mRNA lengths. Additionally, WAGO-1 aberrantly binds small RNAs derived from CSR-1 targets. Cycles of CSR-1 small RNA production are less efficient. CSR-1 target mRNAs maintain protection from WAGO-1 silencing but fail to be fine-tuned.

CSR-1 slicer activity is proposed to slice mRNA targets generating a new 3’ mRNA end that serves as a template for EGO-1 22G-RNA synthesis. Sequential rounds of CSR-1 target slicing and small RNA synthesis by EGO-1 leads to the production of abundant 22G-RNAs derived from the full mRNA length. Our results suggest RHA-1 promotes sequential rounds of 22G-RNA production. Considering RHA-1 is a DExH RNA helicase where its ATPase activity allows processive translocation along RNA from the 3’ to 5’ direction(*32*), we propose RHA-1 may promote mRNA accessibility for CSR-1 or EGO-1 during small RNA biogenesis (Fig. 8). By remodeling these RNA-protein complexes, RHA-1 may facilitate consecutive rounds of CSR-1 mRNA slicing and EGO-1 small RNA synthesis. Alternatively, RHA-1 may promote small RNA loading to CSR-1 while inhibiting loading to WAGO-1 after 22G-RNA synthesis by EGO-1 (Fig. 8). Because WAGO-1 does not possess slicer activity, in the absence of RHA-1, mis-sorted WAGO-1 bound 22G-RNAs could similarly disrupt the amplification of small RNAs derived from the 5’ regions of CSR-1 target mRNAs.

While 22G-RNAs produced from CSR-1 targets are aberrantly sorted to WAGO-1 in *rha-1* mutants, we do not find evidence for gain of WAGO-1-mediated silencing of these targets. Given that we find 3’ CSR-1 22G-RNA targeting is intact in *rha-1* mutants, one model is that the residual level of CSR-1 bound 22G-RNAs is sufficient to protect endogenous protein-coding genes from piRNA and WAGO-1-mediated silencing (Fig. 8). Additionally, these aberrant WAGO-1 22G-RNAs may not be effective in recruiting mRNA degradation factors needed for robust RNA silencing. Since piRNAs and WAGO-1 22G-RNAs normally target about 2000 genes in the germline, 22G-RNA mis-sorting in the absence of RHA-1, results in approximately 3-fold increase to total WAGO target genes. This increase in target genes potentially weakens WAGO-1 recruitment of downstream factors needed for mRNA degradation.

Intriguingly, another conserved RNA helicase ZNFX-1 promotes amplified production of small RNAs at the 3’ ends of WAGO and CSR-1 target mRNAs (*38*, *53*). ZNFX-1 helicase is processive on RNA substrates in the 5’ to 3’ direction, opposite of RHA-1, potentially explaining how these helicases promote small RNA production on the opposite ends of mRNAs (*38*, *54*). This role for ZNFX-1 in balancing small RNA production across target mRNA lengths is proposed to contribute to the stability of epigenetic inheritance across generations (*38*, *53*). Together our findings suggest that in addition to WAGO versus CSR-1 targeting, the site of small RNA production along mRNA target lengths may impact whether a given mRNA is protected or degraded following small RNA targeting. Our study along with others provide valuable insight into the diverse roles for RNA helicases in small RNA-mediated gene regulation.

### Spatial organization of CSR-1 and EGO-1 localization

Another outstanding question in the field is where within germ cells CSR-1 small RNAs are produced and function in gene regulation. While germ granule factors are enriched within specific sub-compartments, these proteins rapidly exchange between granule and cytoplasmic localization (*55*). CSR-1 and EGO-1 localize to the cytoplasm and are also enriched in distinct germ granule sub-compartments, D and E respectively (*43*). Notably, the D compartment is closest to the nucleus just outside of nuclear pores, where CSR-1 is positioned to potentially interact with newly exported mRNAs prior to piRNAs and WAGO-1 located in the P compartment. Because CSR-1 and EGO-1 localize both within germ granules and the cytoplasm, where CSR-1 targeting, slicing, and EGO-1-dependent small RNA production occurs is still under investigation.

In support of EGO-1 small RNA production occurring in the cytoplasm, 22G-RNAs produced from CSR-1 targets exhibit a three-nucleotide periodicity in phase with ribosomal protected fragments (*23*). This suggests either 22G-RNA production or CSR-1 slicing of mRNA targets occurs in the cytoplasm on transcripts undergoing translation. Additionally, the abundance of small RNAs mapping to mRNAs anti-correlates with translation efficiency (*23*). The model follows that efficiently and highly translated mRNAs are inaccessible to both EGO-1 dependent small RNA synthesis and CSR-1 binding, while poorly translated mRNAs with less active ribosomes are more accessible.

Potentially in conflict with this model, specific loss of the E compartment and EGO-1 granule localization by mutations to either ELLI-1 or EGC-1, disrupts EGO-1-dependent small RNA production on 5’ regions of mRNA target lengths (*44*). This suggests EGO-1 must localize to the E compartment at some point in the 22G-RNA production cycle for 22G-RNA synthesis to occur from the 5’ ends of mRNA targets. Taken together, we propose a hybrid model where CSR-1 slicing may first occur in the cytoplasm while coordinated rounds of 22G-RNA synthesis from the 5’ region of mRNAs and/or 22G-RNA loading occurs at the E compartment.

Here we identify RHA-1 as an additional E compartment factor required for CSR-1 22G-RNA production on the 5’ regions of target mRNAs. Unlike ELLI-1 and EGC-1, loss of RHA-1 does not greatly compromise CSR-1 or EGO-1 germ granule formation. Instead, RHA-1 germ granule localization depends on both CSR-1 and EGO-1, suggesting RHA-1 function in CSR-1 small RNA biogenesis and sorting occurs downstream of initial CSR-1 and EGO-1 activity. Additionally, this recruitment may contribute to the specificity of RHA-1 function in CSR-1 small RNA production. Our findings build on the importance of E compartment factors in small RNA production from the 5’ regions of CSR-1 target mRNAs. Together, these data hint at the important coordination between CSR-1 and EGO-1 for small RNA production and mRNA fine-tuning.

### Small RNA roles in male fertility

In *C. elegans*, several Argonaute proteins have been shown to regulate male germ cell differentiation and function, including CSR-1 and PRG-1 (*8*, *56*). In most animals, endo-siRNAs and piRNAs are dynamically produced throughout male germline development and essential for spermatogenesis (*57*, *58*). In male germ cells of mice and fruit flies, groups of endo-siRNAs and piRNAs function to silence endogenous protein coding genes, performing roles outside of genome defense (*9*, *10*). In our study, we find RHA-1 ATPase and CSR-1 slicer activity also contribute to male fertility, likely through the regulation of spermatogenesis-enriched genes. Together, these studies hint at the evolutionary conserved roles of endogenous gene regulation by small RNAs in male germ cell development and function.

Since murine and human orthologs of RHA-1 are essential genes, whether they function similarly in male fertility as we have shown for *C. elegans* remains unknown (*59*, *60*). Intriguingly, in a proteomics study, the RHA-1 murine ortholog DHX9 was found enriched in the chromatoid bodies (CBs) of male germ cells (*61*). CBs are RNA-rich biomolecular condensates, like germ granules, that contain many small RNA pathway and RNA processing factors (*61*). The localization of DHX9 to CBs of male germ cells along with small RNA factors, suggests DHX9 may function similarly in male fertility and small RNA regulation as we show here for RHA-1. Overall, these findings leave open the possibility that RHA-1 performs a conserved role in small RNA regulation within male germ cells. Since multiple small RNA classes are present in the male germ cells of most animals, future work investigating the function of these small RNAs may provide valuable insight into the complex processes required for sperm development and germ cell fertility.

## Methods

### Caenorhabditis elegans strains and maintenance

Animals were grown on standard nematode growth media (NGM) plates seeded with the Escherichia coli OP50 strain at 20 °C or temperatures where indicated. A complete list of strains used in this study are provided in **Supplemental Data 1.**

### Generation of CRISPR-Cas9 lines

#### Cas9/sgRNA constructs

We used the online tool CHOPCHOP to design sgRNAs. The sgRNAs were cloned into pDD162 by overlapping PCR using the plasmid PDD162 (Dickinson et al. 2013) as the PCR template and the appropriate primers with 20 base pair overlap. Overlapping PCR products were inserted into linearized PDD162 vector after SpeI/BsrBI digest by seamless ligation cloning extract (SLiCE) method (*62*).

#### Donor constructs

To generate the GFP::RHA-1 donor construct, 500 bp upstream and 500 bp downstream of the *rha-1* TSS, and the GFP coding sequences were amplified by PCR using N2 genomic DNA or plasmids containing GFP as templates. PCR fragments were inserted into linearized pUC19 digested with HindIII/KpnI and ligated by SLiCE. To generate amino acid substitutions for RHA-1(K410Q) and RHA-1(E505A), single-strand oligodeoxynucleotides (ssODNs) were used as donor templates designed with the single amino acid substitution, silent mutations to the sgRNA site to prevent recutting, and silent mutations to insert new restriction enzyme cut site for easy genotyping.

#### Cas9 mediated large gene deletions

To generate *elli-1* mutant alleles *(uoc35)* and *(uoc36)* two crRNAs approximately 1000 nucleotides apart with the 5’ crRNA targeting near the start codon were injected at the same time following the protocol detailed in (*63*).

### RNA interference

RNAi was performed by feeding animals with *E. coli* HT115 (DE3) strains expressing the appropriate dsRNA. RNAi bacterial strains were obtained from the Ahringer *C. elegans* RNAi Collection (Source BioScience). Bacterial cultures were grown in Luria broth supplemented with 100 μg/mL ampicillin and 50 μg/mL tetracycline overnight at 37°C. Cultures were seeded on NGM plates containing 100 μg/mL ampicillin, 50 μg/mL tetracycline, and 1 mM IPTG and incubated at room temperature for 24 hours. HT115 (DE3) expressing empty RNAi vector L4440 was used for control.

#### RNAi of germline genes

For RNAi of *pos-1* and *mex-3* genes, five L1 worms were moved to RNAi and control L4440 plates and performed in triplicate. Three days later, adult worms were removed from plates. The following day unhatched embryos and hatched progeny were counted for each plate. Plots show percentage of unhatched embryos over sum of unhatched embryos and hatched progeny.

#### RNAi of somatic genes

For RNAi of *unc-22* and *dpy-13* genes, twenty-five L1 worms were transferred to RNAi and control L4440 plates, performed in triplicate. Three days later, adult worms were scored for phenotypes. For RNAi of *unc-22*, worms were scored for body muscle twitching. For RNAi of *dpy-13*, worms were scored for a dumpy and shortened body.

#### RNAi for RHA-1 granule counts

For RHA-1 granule counts, twenty L1 worms were transferred to RNAi and control L4440 plates, performed in triplicate. Three days later, the germlines of young adult animals were imaged following the confocal live imaging method (see details below). Number of GFP::RHA-1 granules were counted per germ cell nucleus as distinct puncta localized outside of mCh::NPP-7 signal at the nuclear periphery.

### Mating assay with auxin-induced CSR-1 degradation

The degron-tagged CSR-1 strains (no rescue, transgenic rescue of wild type CSR-1, and transgenic rescue of catalytic dead CSR-1) were self-crossed with males of the same genotype to maintain males in their population. Populations of L1 worms were transferred to NGM plates containing 500 µM auxin, 0.5% ethanol plates or 0.5% ethanol control plates. After 48 h, male worms were transferred to NGM cross plates containing 500 µM auxin, 0.5% ethanol plates or 0.5% ethanol control plates with obligate *fog-2* females. After 48h of mating, single *fog-2* females were transferred to NGM plates and hatched progeny was counted for brood size. For strains grown on auxin, CSR-1 depletion was confirmed by live imaging. Transgenic rescue strains were confirmed for transgene expression by live fluorescence imaging.

### Brood size and mating assay

#### Brood size

Individual L4 worms were transferred to fresh NGM plates and grown at 20 °C. Worms were transferred daily to fresh NGM plates to allow for easy quantification of progeny until no more eggs were laid. Progeny were counted 48 hours after transferring the parent and number of progeny per individual hermaphrodite was totaled.

#### Germline mortality assay

First, *rha-1(tm329)* mutants and RHA-1 ATPase mutants were outcrossed two times with wild type males to generate newly homozygous mutant strains. Ten adult worms for these mutant strains along with wild type or control *gfp::rha-1* strains were moved to 25°C designated as the P0 generation. Ten L3 progeny from these worms were transferred to a new plate designated the F1 generation. This was performed for each subsequent generation. To test fertility of a given generation, twelve L3 worms for each generation were singled to new plates for brood size measurements.

#### Mating assay

For 20 °C mating assay, L3 males were isolated and grown at 20 °C. The next day single males were transferred to cross plate with single L4 obligate females and grown at 20 °C. After 24h, individual females were transferred to fresh NGM plate and resulting brood size was measured as described above. For 25 °C mating assay, L3 males were isolated and grown at 25 °C. The next day six males were transferred to cross plate with six L4 obligate females and grown at 25 °C. The increased number of worms per mating plate here was to ensure worms mated as we found reduced male mating behavior at 25 °C for both wild type and mutant males. After 24h, individual females were transferred to fresh NGM plate and resulting brood size was measured as described above.

### Sperm activation assay

L4 males were isolated and kept overnight at 25 °C for 24 hours. Spermatids were isolated from 12 males washed and dissected on slides in sperm medium (50mM HEPES pH7.8, 50mM NaCl, 25mM KCl, 5mM CaCl_2_, and 1mM MgSO_4_) activated with 20μg/mL pronase E. Spermatids were imaged 30 min after dissection and scored by three categories: spermatids with normal pseudopods (wild type), spermatids with irregular pseudopods (irregular), and round spermatids with no pseudopods (inactivated).

### Confocal live imaging

Tagged fluorescent proteins were visualized in living nematodes by mounting young adult animals on 2% agarose pads with M9 buffer (22 mM KH_2_PO4, 42 mM Na_2_HPO4, and 86 mM NaCl) with 20 mM levamisole. Fluorescent images used for localization studies were captured using a Zeiss LSM800 confocal microscope with a Plan-Apochromat 40×/1.4 Oil objective.

### Quantitative real-time PCR

1 μg of total RNA was reverse transcribed with SuperScript IV Reverse Transcriptase (Invitrogen) in 1 × reaction buffer, 2 U SUPERase-In RNase Inhibitor (Invitrogen), 0.5 mM dNTPs, and 2.5 μM random hexamers. Each real-time PCR reaction consisted of 3 μL of cDNA, 1 μM forward gene-specific primer and 1 μM reverse gene-specific primer. The amplification was performed using iTaq Universal SYBR Green Supermix (Bio-Rad) on the Bio-Rad CFX96 Touch Real-Time PCR Detection System. The experiments were repeated for a total of three technical replicates. tba-1 expression was used as a housekeeping gene.

### Total small RNA-seq library preparation

Total RNA was extracted using the standard method with TRIzol reagent (Invitrogen) from whole animals of ∼100,000 synchronized young adults grown at 20 °C. Small (<200 nt) RNAs were enriched with mirVana miRNA Isolation Kit (Ambion). In brief, 80 μL (200–300 μg) of total RNA, 400 μL of mirVana lysis/binding buffer and 48 μL of mirVana homogenate buffer were mixed well and incubated at room temperature for 5 min. Then, 176 μL of 100% ethanol was added and samples were spun at 2500×g for 4 min at room temperature to pellet large (>200 nt) RNAs. The supernatant was transferred to a new tube and small (<200 nt) RNAs were precipitated with pre-cooled isopropanol at −80 °C. Small RNAs were pelleted at 20,000×g at 4 °C for 30 min, washed once with 70% pre-cooled ethanol, and dissolved with nuclease-free water. Ten micrograms of small RNAs were fractionated on a 15% PAGE/7M urea gel, and RNA from 17 nt to 40 nt was excised from the gel. RNA was extracted by soaking the gel in 2 gel volumes of NaCl TE buffer (0.3 M NaCl, 10 mM Tris-HCl, 1 mM EDTA pH 7.5) overnight. The supernatant was collected through a gel filtration column. RNA was precipitated with isopropanol, washed once with 70% ethanol, and resuspended with 15 μL nuclease-free water. RNA samples were treated with RppH to convert 22G-RNA 5′ triphosphates to monophosphates in 1 × reaction buffer, 10 U RppH (New England Biolabs), and 20 U SUPERase-In RNase Inhibitor (Invitrogen) for 3 h at 37 °C, followed by 5 min at 65 °C to inactivate RppH. RNA was then concentrated with the RNA Clean and Concentrator-5 Kit (Zymo Research). Small RNA libraries were prepared according to the manufacturer’s protocol of the NEBNext Multiplex Small RNA Sample Prep Set for Illumina-Library Preparation (New England Biolabs). NEBNext Multiplex Oligos for Illumina Index Primers were used for library preparation (New England Biolabs). Libraries were sequenced using an Illumina HiSeq4000 to obtain single-end 50 nt sequences at the University of Chicago Genomic Facility.

### Immunoprecipitation – small RNA-seq

A total of 100,000 synchronized young adult animals were used for immunoprecipitation. Synchronized animals were washed with M9 buffer three times before frozen in liquid nitrogen and stored at −80 °C. Worm pellets were resuspended in equal volumes of immunoprecipitation buffer (20 mM Tris-HCl pH 7.5, 150 mM NaCl, 2.5 mM MgCl_2_, 0.5% NP-40, 80 U/mL RNase Inhibitor (Thermo Fisher Scientific), 1 mM dithiothreitol, and protease inhibitor cocktail without EDTA (Promega)), and grinded in a stainless steel dounce tissue grinder until cuticles of most worms were broken down. Lysates were clarified by spinning down at 15,000 rpm, 4 °C, for 15 min. Supernatants were incubated with the GFP-Trap magnetic agarose beads (ChromoTek) at 4 °C for 1 h. Beads were washed with wash buffer (20 mM Tris-HCl pH 7.5, 150 mM NaCl, 2.5 mM MgCl_2_, 0.5% NP-40, and 1 mM dithiothreitol) six times, and then resuspended in TBS buffer for RNA extraction. Total RNA was extracted using the standard method with TRIzol reagent (Invitrogen). Small RNA libraries were prepared as described above and sequenced using Illumina NovaSeq 6000 to obtain single-end 50 nt sequences at the University of Chicago Genomic Facility.

### Poly A RNA-seq library preparation

Total RNA was extracted using the standard method with TRIzol reagent (Invitrogen) from whole animals of ∼100,000 synchronized young adults. The Promega PolyATtract mRNA Isolation Systems was used to isolate the polyA mRNA fraction following the small-scale isolation protocol provided by Promega, starting with 100 µg total RNA. Next, RNA libraries were prepared with NEBNext Ultra II Directional RNA Library Prep Kit for Illumina (New England Biolabs) starting with 10ng of total polyA mRNA per sample. NEBNext Multiplex Oligos for Illumina Index Primers were used for library preparation (New England Biolabs). RNA libraries were sequenced using an Illumina NovaSeq to obtain paired-end 50 nt sequences at the University of Chicago Genomic Facility.

### Single-molecule fluorescent in-situ hybridization (smFISH)

Whole worm smFISH was performed on staged L4 or young adult worms. Two technical replicates were performed for each genotype and condition. 100 staged worms were picked into M9 buffer in 1.5mL microcentrifuge tubes and washed with M9 three times. Worms were then fixed in 4% formaldehyde for 45 min at room temperature. Samples were then washed with PBS and dehydrated with 70% ethanol at 4 °C for two overnights. Gonads were re-hydrated with PBS and washed once with FISH wash buffer (2× SSC, 10% formamide) at 37 °C. FISH probes were suspended in Hybridization Buffer (10% formamide, 2 mM vanadyl ribonucleoside complex, 20 mg/mL BSA, 10 mg/mL dextran sulfate, 2 mg/mL *E. coli* tRNA) 1:50. smFISH probes were created by Biosearch Technologies to target *klp-18* mRNA or *gipc-1* mRNA. The *klp-18* mRNA and *gipc-1* mRNA probes were conjugated to Quasar670 (Custom probe, Biosearch Technologies Cat. No. SMF-1065-5). Samples were hybridized overnight at 37 °C in dark. Samples were washed with FISH wash buffer, stained with DAPI, and mounted onto 25 mm × 75 mm × 1 mm plain glass slides with ProLong Diamond Antifade Mountant. Samples were imaged on a Zeiss LSM800 Confocal Microscope at ×40 magnification using a Plan-Apochromat ×40/1.4 Oil Objective.

### Image processing and analysis

Image processing and quantification of fluorescent puncta were performed using ImageJ. Single slice images of gonads were used for quantification. ROIs were selected, and areas of ROIs were measured. For measurement of granule sizes, the same threshold and lower limit was applied across all germline images for control and mutants to eliminate background signal. Then the area in µm^2^ was measured per granule. Granule area was measured for at least 10 gonads with the same ROI of about 10-12 pachytene nuclei per gonad. For quantitative co-localization analysis, all image manipulations were performed with ImageJ using the Coloc 2 plugin. Images of 8–10 gonads were collected and quantified. For smFISH (Quasar670), a rolling ball background subtraction was used across all images set manually and uniformly across all samples. Images from 20–30 gonads were collected. Signal from the RNA probes was quantified by measuring mean fluorescent intensity in the same ROI for each gonad. Data were compared by two-tailed p values calculated using Mann-Whitney-Wilcoxon test.

### Sequencing data analysis

#### RNA-seq

Fastq reads were trimmed of adaptors using cutadapt (*64*). Trimmed reads were aligned to the C.elegans genome build WS230 using bowtie2 ver 2.3.0 (*65*). After alignment, reads were overlapped with genomic features (protein-coding genes, pseudogenes, transposons) using bedtools intersect (*66*). Reads per kilobase million (RPKM) values were then calculated for each individual feature by summing the total reads mapping to that feature, multiplied by 1e6, and divided by the product of the kilobase length of the feature and the total number of reads mapping to protein-coding genes. Protein-coding genes were used to normalize by sequencing depth because mRNA libraries were prepared by polyA tail selection. RPKM values were then used in all downstream analyses using custom R scripts, which rely on packages ggplot2, reshape2, ggpubr, dplyr (*67*). Differential expression analysis was performed using DESeq2 with an adjusted *p* value ≤0.05 (*68*).

#### Small RNA-seq

Fastq reads were trimmed using custom perl scripts. Trimmed reads were aligned to the *C. elegans* genome build WS230 using bowtie ver 1.2.1.1 with options -v 0 -best -strata (*69*). After alignment, reads that were between 17–40 nucleotides in length were overlapped with genomic features (rRNAs, tRNAs, snoRNAs, miRNAs, piRNAs, protein-coding genes, pseudogenes, transposons) using bedtools intersect (*66*). Sense and antisense read mapping to individual miRNAs, piRNAs, protein-coding genes, pseudogenes, RNA/DNA transposons, simple repeats, and satellites were totaled and normalized to reads per million (RPM) by multiplying be 1e6 and dividing read counts by total mapped reads, minus reads mapping to structural RNAs (rRNAs, tRNAs, snoRNAs). Reads mapping to multiple loci were penalized by dividing the read count by the number of loci they perfectly aligned to. Reads mapping to miRNAs and piRNAs were only considered if they matched the sense annotation without any overlap. 22G-RNAs were defined as 21 to 23 nucleotide long reads with a 5′G that aligned to protein-coding genes, pseudogenes, or transposons. RPM values were then used in all downstream analyses using custom R scripts using R version 4.0.0, which rely on packages ggplot2, reshape2, ggpubr, dplyr.

#### Metagene analysis

Metagene profiles across gene lengths were calculated by computing the depth at each genomic position using 21–23 nucleotide long small RNA reads with a 5′G using bedtools genomecov(*66*). A custom R script was then used to divide genes into 100 bins and sum the normalized depth within each bin. Groups of genes were then plotted using the sum of the normalized depth at each bin with units in reads per million (RPM). The read distribution metagene normalizes the RPM metagene by dividing the sum of reads for each gene per bin by the total read count of the entire gene. From this, every gene contributes one unit so the area under each trace is the same between the compared samples. The metagene for IP-small RNA-seq data presents the read count mean plus or minus one standard deviation plotted for groups of gene sets.

## Supporting information

Supplementary infomation

## Acknowledgments

Some strains used in this study were provided by the Caenorhabditis Genetics Center (CGC) funded by NIH Office of Research Infrastructure Programs (P40 OD010440).

## Funding

This work is supported in part by NIH predoctoral training grant (T32 GM007183) to O.G.; the NIH grant R01-GM132457 to H.-C.L.

## Author contributions

Conceptualization: O.G. and H.-C.L., Investigation and data analyses: O.G., Bioinformatics analyses: J.B. and W.-S.W., Writing: O.G. and H.-C.L.

## Competing interests

The authors declare that they have no competing interests.

## Notes

### Competing Interest Statement

The authors have declared no competing interest.

