## Supplementary infomation for "RNA Helicase A promotes small RNA biogenesis and sorting in germ cells"

**Title:**

**This file includes:**

**9 Supplementary Figures**

**1 Supplementary Data File**

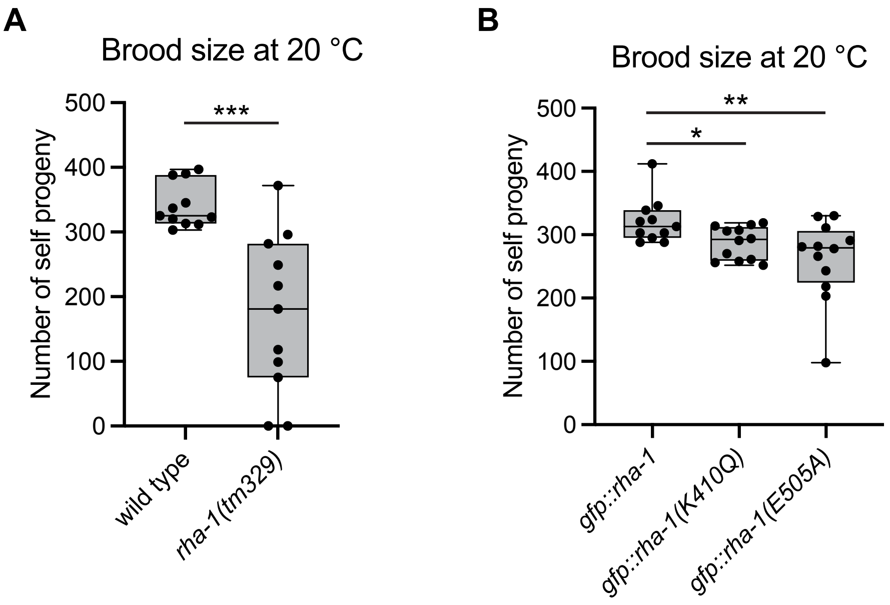


**Figure S1. RHA-1 ATPase activity is required for optimal fertility.**

(**A** and **B**) Brood sizes of the indicated animals grown at 20 °C. *N* = 11-12 animals. For the boxplots, the line indicates the median value, the box indicates the first and third quartiles, and the whiskers indicate the min and max values. Two-tailed p values were calculated using Mann-Whitney-Wilcoxon test. *, $p<0.05$; **, $p<0.01$; ***, $p<0.001$.

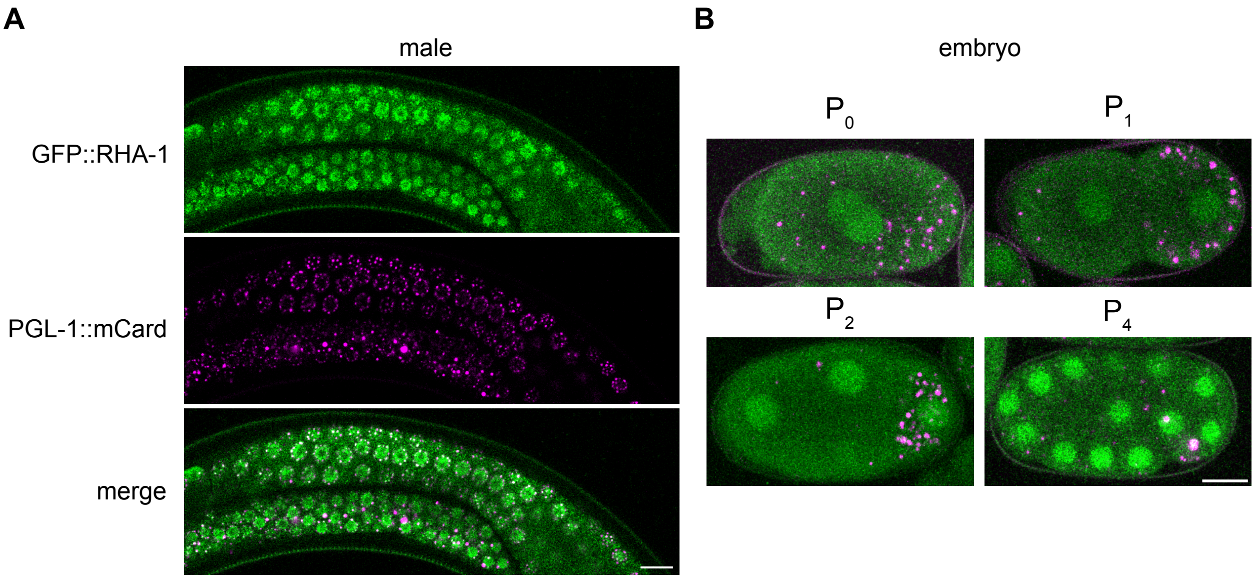


**Figure S2. RHA-1 is expressed in the male germline and developing embryos.**

(**A**) Representative live-fluorescent images of animals expressing GFP::RHA-1 and PGL-1::mCardinal (P compartment marker) in the young adult male germline. (**B**) Representative live-fluorescent images of animals expressing GFP::RHA-1 and PGL-1::mCardinal (P compartment marker) in the four labeled stages of developing embryos. Scale bars = 10 μm.

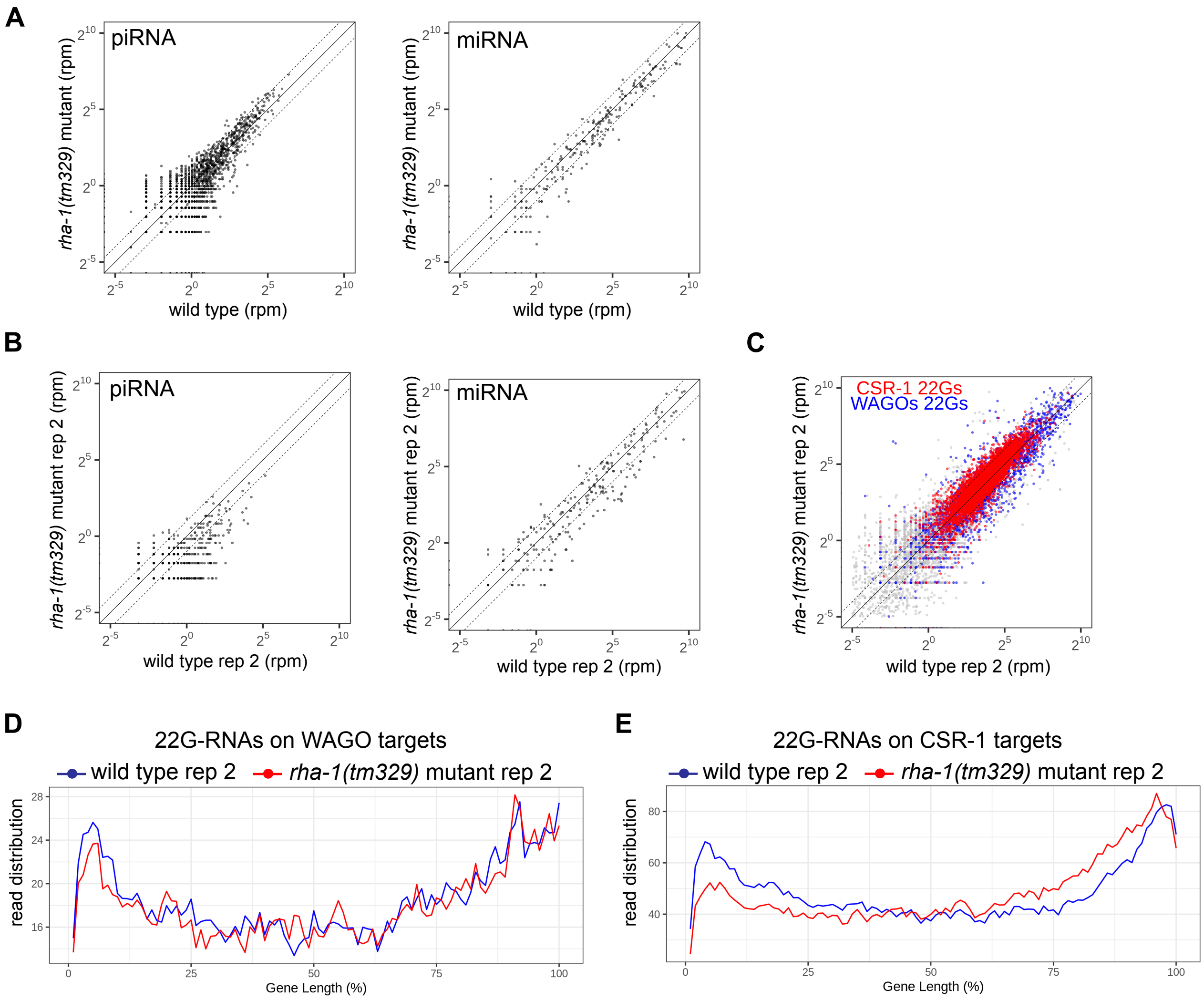


**Figure S3. Total small RNA abundance is relatively unchanged in *rha-1* mutants.**

(**A**) Scatterplots showing the abundance of piRNAs (left) or miRNAs (right) in wild type worms compared to *rha-1(tm329)* mutant worms for replicate 1. (**B**) Scatterplots showing the abundance of piRNAs (left), miRNAs (middle), or all 22G-RNAs (right) mapped to each CSR-1 target (red) and WAGO target (blue) in wild type worms compared to *rha-1(tm329)* mutant worms for replicate 2. (**C** and **D**) Metagene traces show the distribution of normalized 22G-RNAs (sRNA-seq) in arbitrary units mapping to WAGO (**C**) or CSR-1 (**D**) target genes by percentage of WAGO or CSR-1 target gene length. Reads from wild type (blue) and *rha-1(tm329)* mutant (red) for replicate 2. For all scatterplots the three diagonal lines indicate a two-fold increase (top), no change (middle), or a two-fold depletion (bottom) in indicated *rha-1* mutant animals.

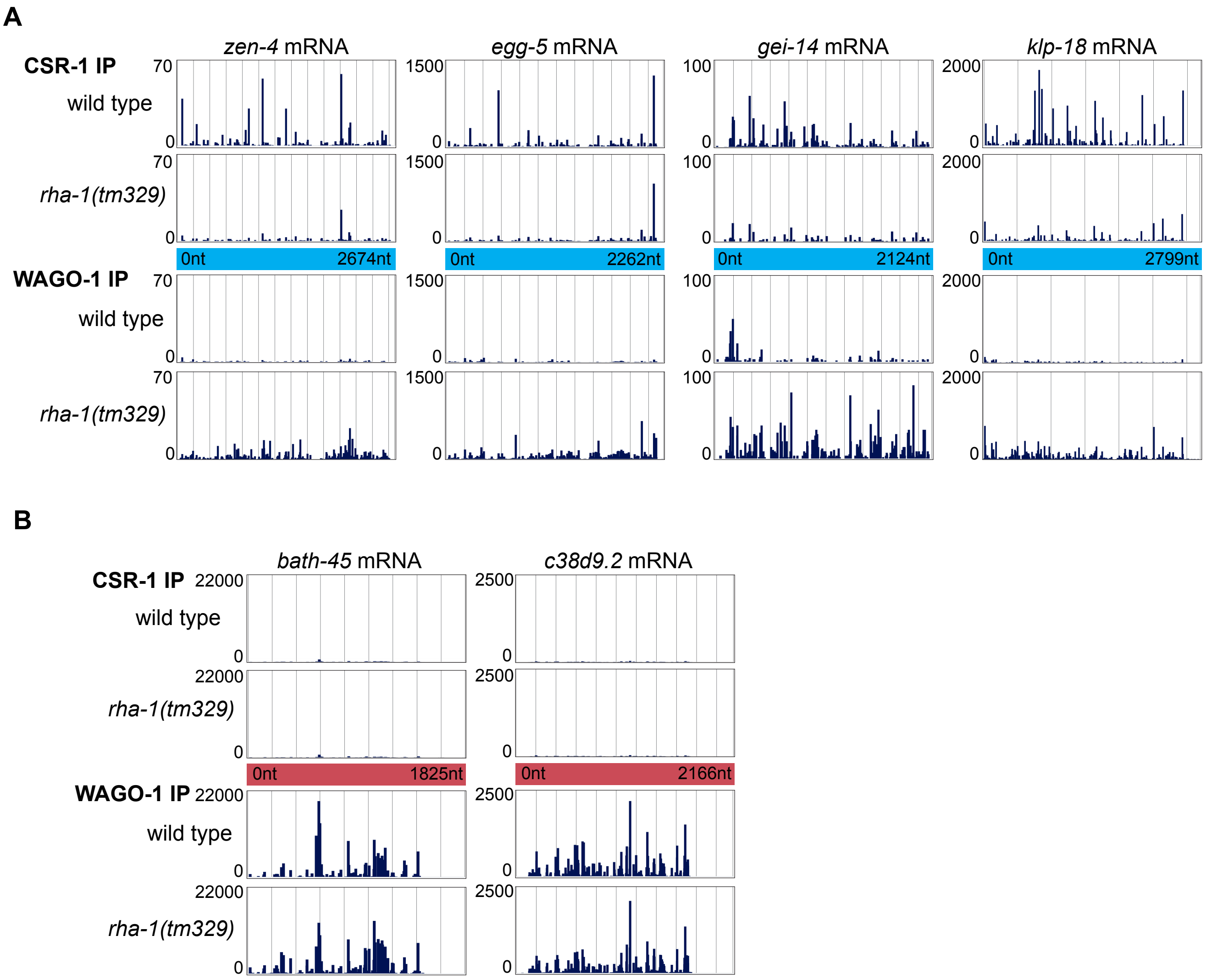


**Figure S4. RHA-1 contributes to the generation of CSR-1 bound 22G-RNAs mapping to the 5' region of CSR-1 target mRNAs.**

(**A** and **B**) Examples of normalized antisense 22G-RNAs bound by either CSR-1 or WAGO-1 in the wild type or *rha-1(tm329)* mutant samples mapping to the indicated CSR-1 target mRNAs (**A**) or WAGO-1 target mRNAs (**B**).

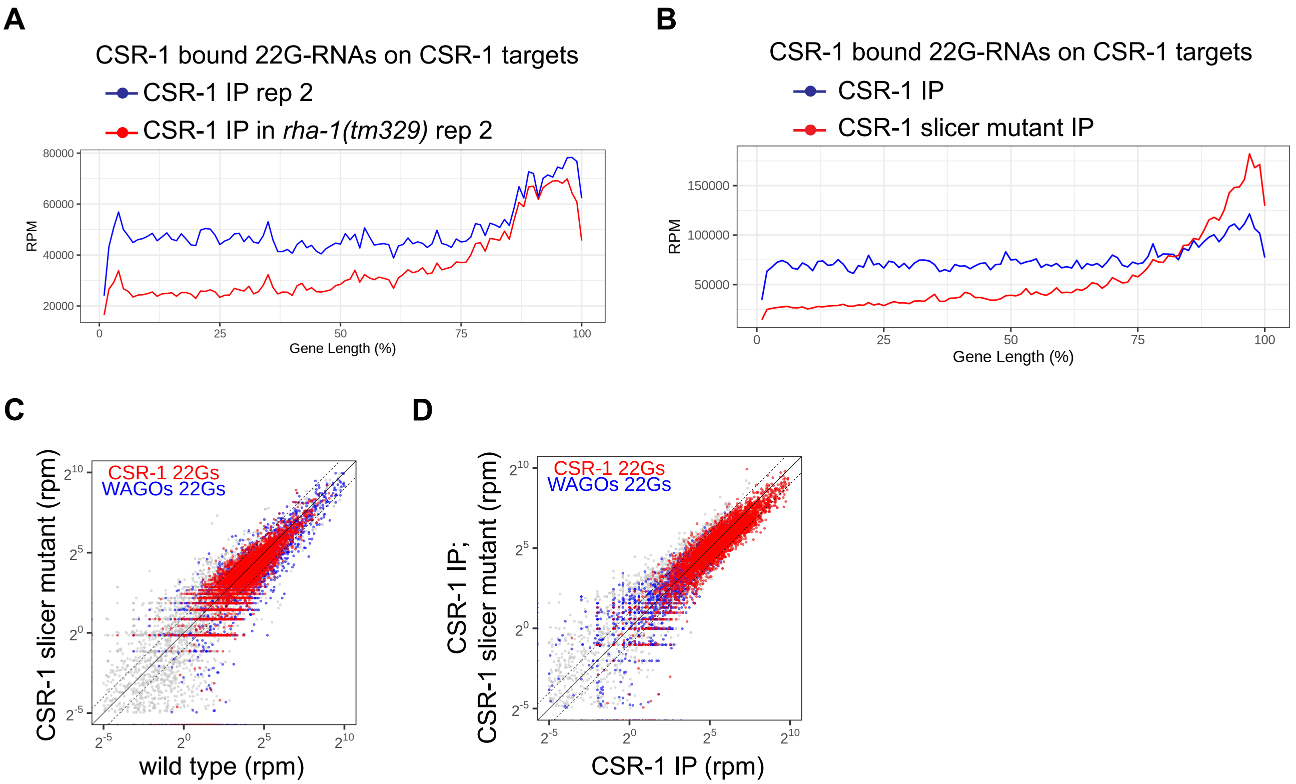


**Figure S5. RHA-1 and CSR-1 slicer activity promote 22G-RNA production from 5' region of mRNAs.**

(**A**) Metagene traces show the distribution of normalized CSR-1 bound 22G-RNAs (IP-sRNA-seq) in reads per million (RPM) mapping to CSR-1 target genes by percentage of CSR-1 target gene length. Reads from wild type (blue) and *rha-1(tm329)* mutant (red). (**B**) Metagene traces show the distribution of normalized CSR-1 bound 22G-RNAs (IP-sRNA-seq) in reads per million (RPM) mapping to CSR-1 target genes by percentage of CSR-1 target gene length. Reads from wild type (blue) and CSR-1 slicer mutant (red). (**C** and **D**) Scatterplots showing the abundance of all 22G-RNAs mapped to each CSR-1 target (red) and WAGO target (blue) in wild type worms compared to CSR-1 slicer mutant worms from total small RNAs (sRNA-seq) (**C**) or CSR-1 bound 22G-RNAs (IP-sRNA-seq) (**D**). The three diagonal lines indicate a two-fold increase (top), no change (middle), or a two-fold depletion (bottom) in CSR-1 slicer mutant animals. (**B** to **D**) Data re-analyzed from published data sets in (Singh et *al.*, 2021).

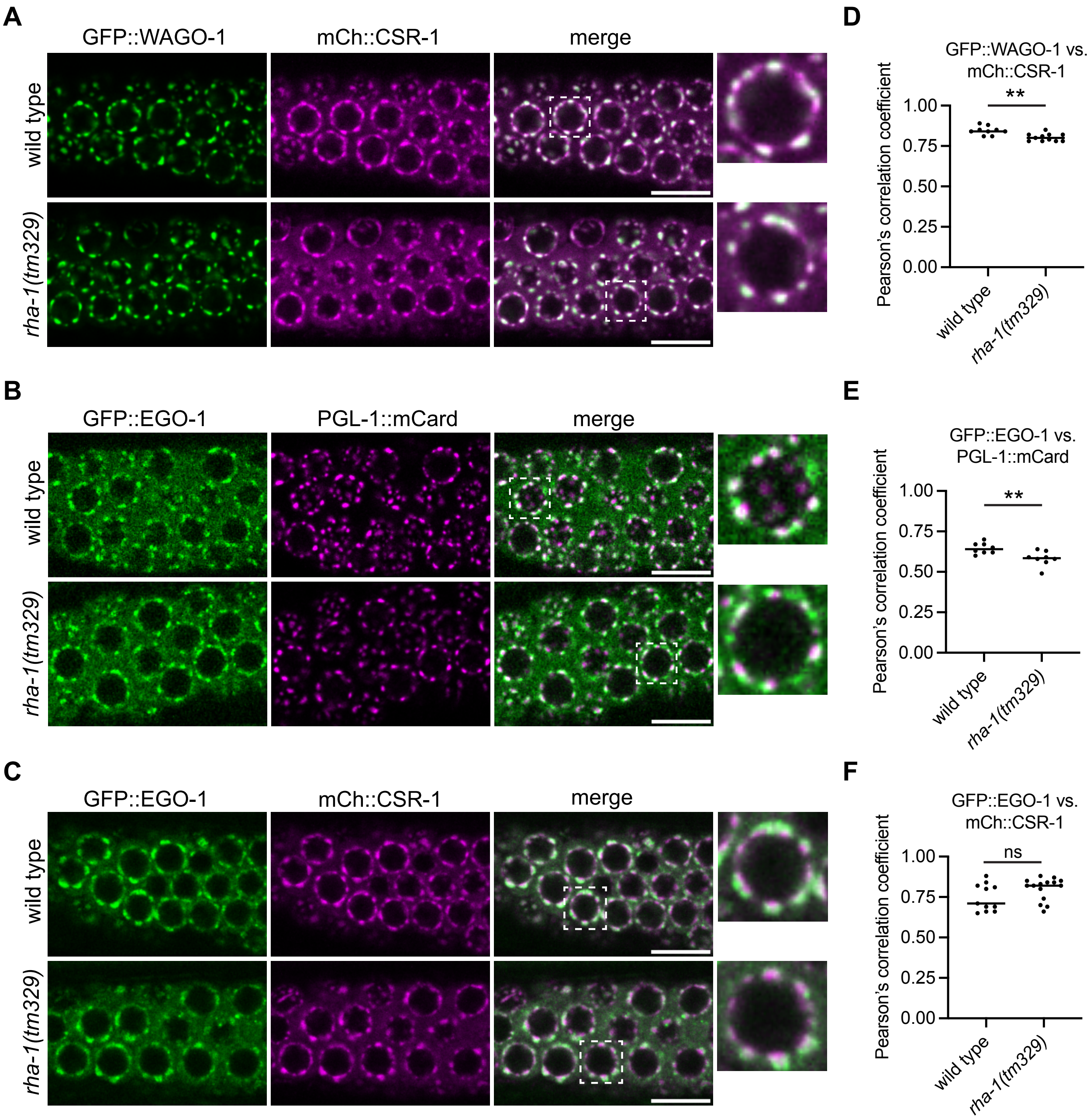


**Figure S6. The P compartment shows reduced co-localization with D and E compartments in *rha-1* mutants.**

(**A** to **C**) Representative live-fluorescent images of pachytene germ cells show the localization of the indicated germ granule markers in wild type and *rha-1(tm329)* mutant animals. Co-marked strains for comparison of the P compartment (GFP::WAGO-1) to the D compartment (mCh::CSR-1) (**A**) or E compartment (GFP::EGO-1) to the P compartment (PGL-1::mCard) (**B**) or E compartment (GFP::EGO-1) to the D compartment (mCh::CSR-1) (**C**). For all images boxes with dashed outline show the single cropped and enlarged nucleus. Scale bars = 10 µm. (**D** to **F**) Quantification of co-localization between the indicated fluorescent proteins of pachytene germ cells. Each data point represents the Pearson's R-value showing the degree of colocalization between two fluorescence channels indicated. The solid black line indicates the mean value. Each data point represents independent gonad images. *N* = 8-15 gonads. Two-tailed p values were calculated using Mann-Whitney-Wilcoxon test. **, $p<0.01$.

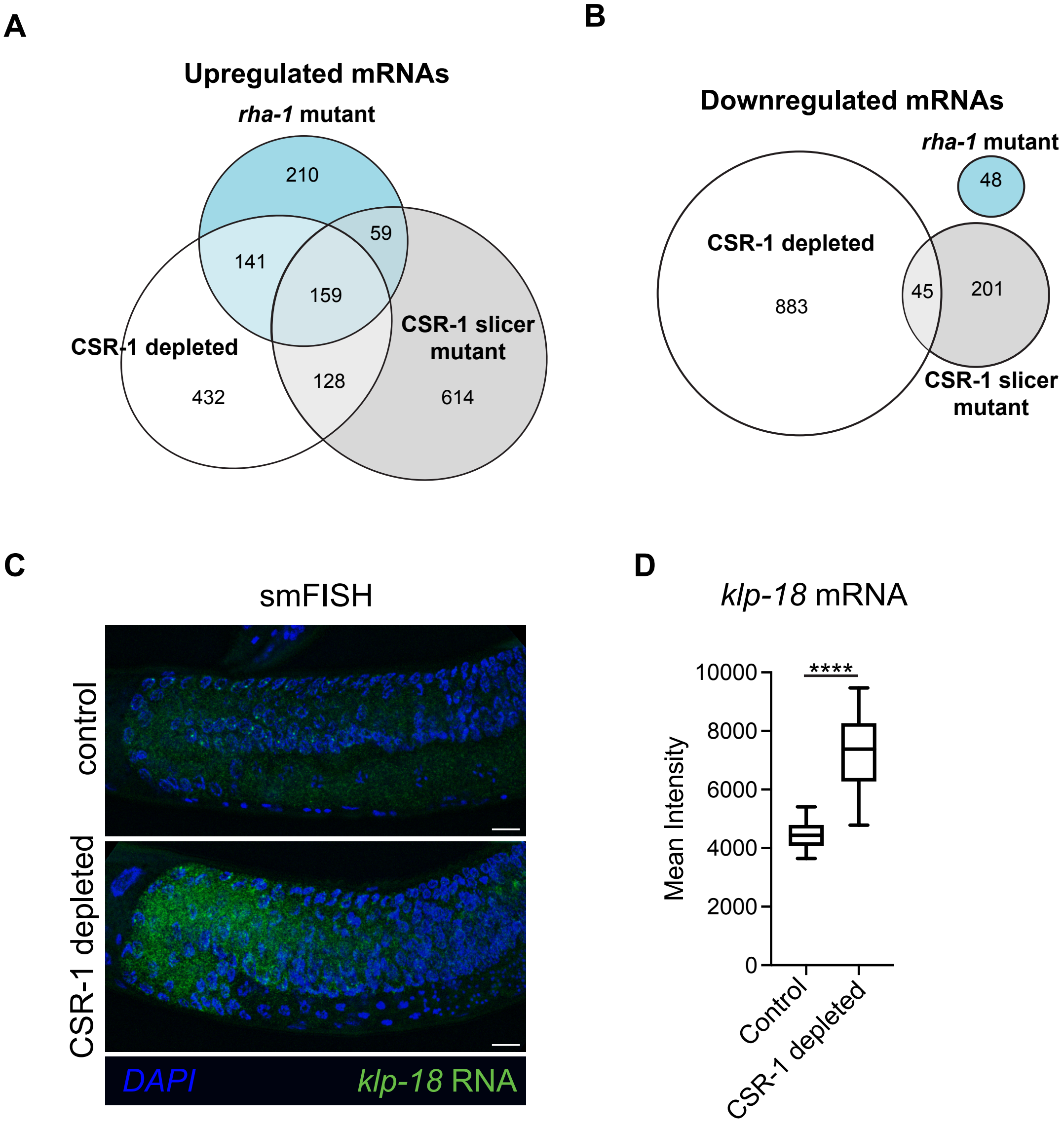


**Figure S7. Loss of RHA-1 results in similar mRNA expression changes as CSR-1 compromised animals.**

(**A** and **B**) Proportional Venn diagrams showing the overlap in number of shared significantly upregulated (**A**) or downregulated (**B**) mRNAs in *rha-1(329)* mutant (blue) CSR-1 depleted (white) and CSR-1 slicer mutant (gray) animals. The differentially regulated genes were computed from two biological replicates using DESeq2 adjusted p value of ≤0.05. Data was re-analyzed from published mRNA-seq data sets for CSR-1 depleted (Shen et *al.*, 2018) and CSR-1 slicer mutants (Gerson-Gurwitz et *al.*, 2016). (**C**) Representative images of *klp-18* mRNA smFISH in adult germlines of control (no auxin) and CSR-1 depleted by auxin-induced degradation. DAPI stain (blue) and *klp-18* mRNA (green). Scale bars = 10 µm. (**D**) Boxplot depicting smFISH quantification of *klp-18* mRNA in control and CSR-1 depleted adult germlines. *N* = 10-12 independent animals. For the boxplot, the line indicates the median value, the box indicates the first and third quartiles, and the whiskers indicate the minimum and maximum. Two-tailed p values were calculated using Mann-Whitney-Wilcoxon test. ****, $p<0.0001$.

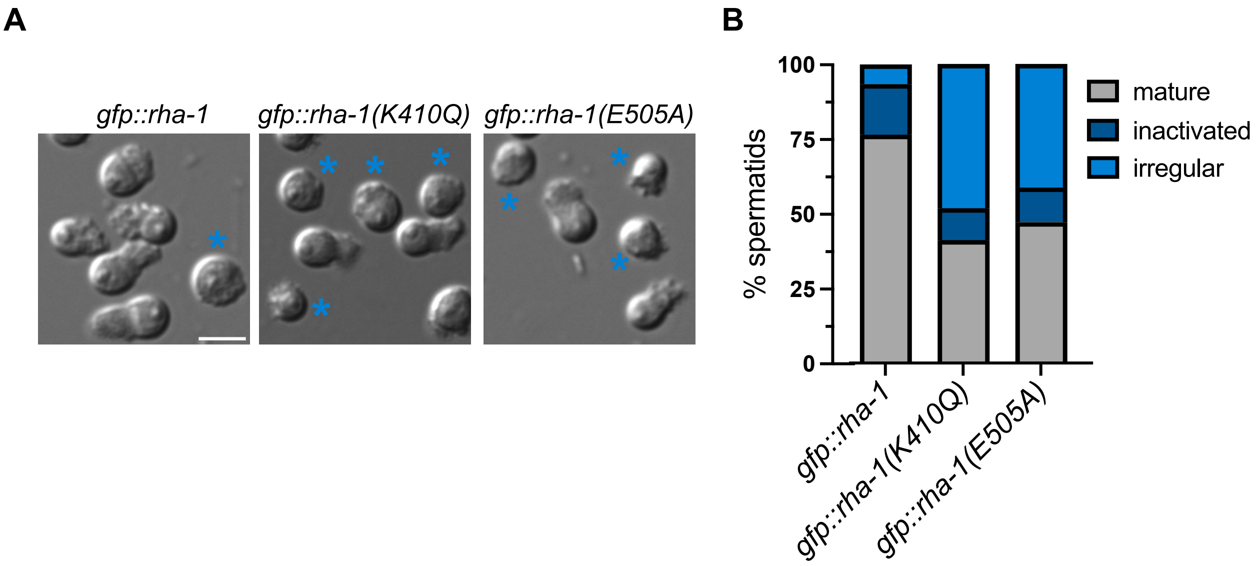


**Figure S8. RHA-1 ATPase activity is required for optimal sperm differentiation.**

(**A**) Representative images of *in vitro* activated sperm by pronase treatment from the indicated strains. Light blue asterisk mark irregular sperm. Scale bar = 5 μm. (**B**) Stacked bar graph showing percentage of activated mature, inactivated, and irregular spermatids from *in vitro* sperm activation assay for the indicated strains. Ten adult male animals were dissected for each strain. More than 1000 spermatids were scored per strain.

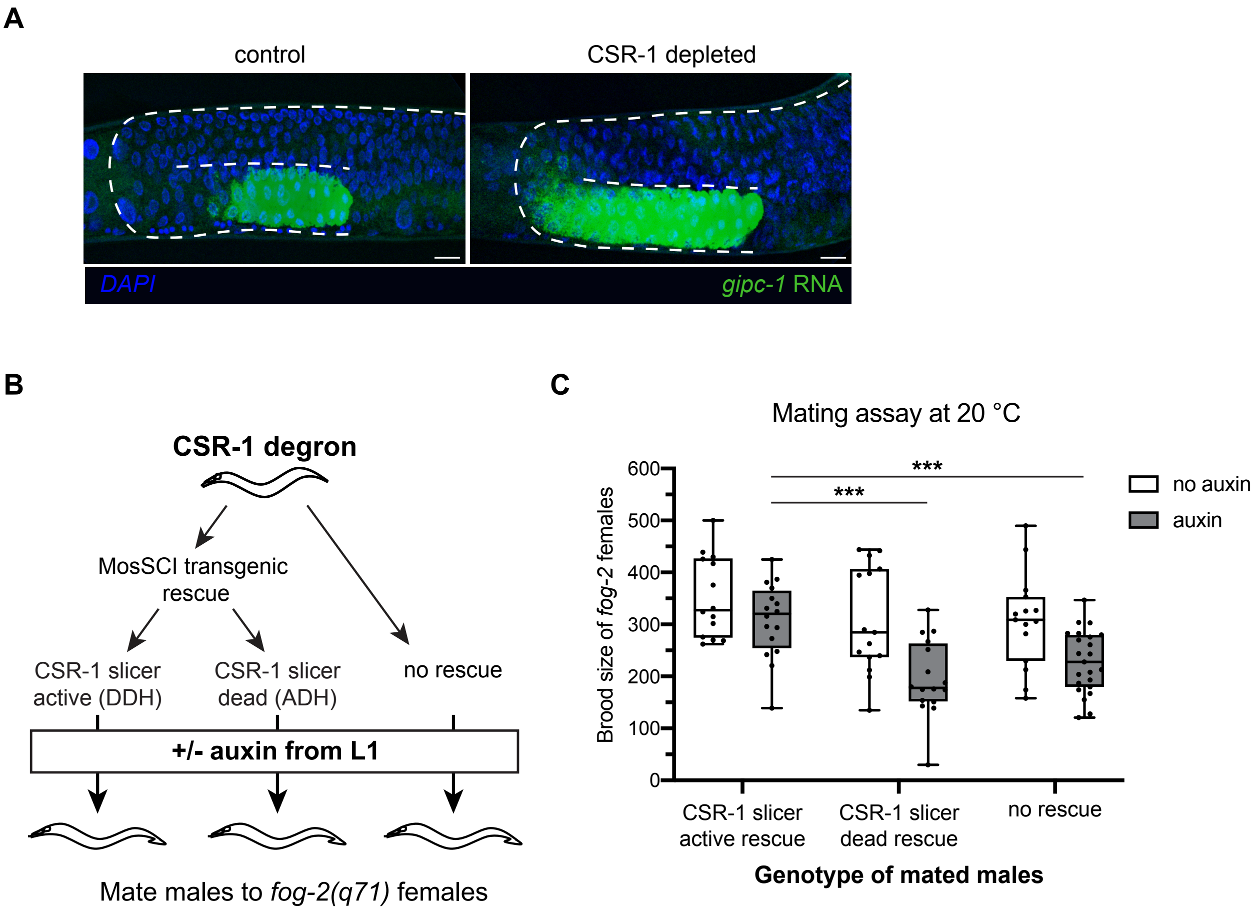


**Figure S9. CSR-1 slicer activity is required for optimal male fertility.**

(**A**) Representative images of *gipc-1* mRNA smFISH in L4 stage control and CSR-1 depleted animals. DAPI stain (blue) and *gipc-1* mRNA (green). Dashed line marks the border of the germline. Scale bar = 10 μm. (**B**) Schematic showing the strains and experimental setup for male mating assay. (**C**) Brood sizes from *fog-2(q71)* females mated to males of the indicated strains grown without auxin (white boxplots) as controls or with auxin (gray boxplots) to deplete endogenous CSR-1. *N* = 14-20 independent animals mated. For the boxplots, the line indicates the median value, the box indicates the first and third quartiles, and the whiskers indicate the minimum and maximum. Two-tailed p values were calculated using Mann-Whitney-Wilcoxon test. ***, $p<0.001$.

**Supplementary Data 1. List of strains**

| Strain | Genotype | Gene(s) | Description | Origin |
|---|---|---|---|---|
| N2 | *C. elegans wild isolate (Bristol)* | | Reference C. elegans. Variety Bristol. | CGC |
| KMW1 | *haf-6(ne335) I; rha-1(tm329) II* | *haf-6; rha-1* | | CGC |
| HCL226 | *rha-1(tm329) II* | *rha-1* | KMW1 strain from CGC outcrossed with N2 | This study |
| HCL207 | *rha-1(uoc24[gfp::rha-1]) II* | *rha-1* | GFP knockin by CRISPR-Cas9 to RHA-1 transcription start site at endogenous locus | This study |
| HCL209 | *rha-1(uoc25[gfp::rha-1[K410Q]]) II* | *rha-1* | GFP::RHA-1 CRISPR-Cas9 amino acid substitution K410Q for ATP binding mutant | This study |
| HCL210 | *rha-1(uoc26[gfp::rha-1[E505A]]) II* | *rha-1* | GFP::RHA-1 CRISPR-Cas9 amino acid substitution E505A for ATP hydrolysis mutant | This study |
| HCL105 | *csr-1(uoc[gfp::csr-1]) IV* | *csr-1* | GFP knockin by CRISPR-Cas9 at start site of CSR-1 isoform b, tags both CSR-1A and CSR-1B isoforms | W Chen et al. 2020 |
| JMC219 | *tor113[GFP::3xFLAG::wago-1] I* | *wago-1* | | CGC |
| HCL378 | *tor113[GFP::3xFLAG::wago-1] I; rha-1(tm329) II* | *wago-1; rha-1* | JMC219 crossed to HCL226 | This study |
| SHG1936 | *ego-1(ust356[mCherry::ego-1]) I* | *ego-1* | | CGC |
| HCL384 | *ego-1(ust356[mCherry::ego-1]) I; rha-1(uoc24[gfp::rha-1]) II* | *ego-1; rha-1* | HCL207 crossed to SHG1936 | This study |
| SHG1941 | *ustIS268[mex-5p::tagrfp::elli-1::tbb-2_3'UTR] I* | *elli-1* | | X Chen et al. 2024 |
| HCL386 | *ustIS268[mex-5p::tagrfp::elli-1::tbb-2_3'UTR] I; rha-1(uoc24[gfp::rha-1]) II* | *elli-1; rha-1* | HCL207 crossed to SHG1941 | This study |
| HCL387 | *ustIS268[mex-5p::tagrfp::elli-1::tbb-2_3'UTR] I; rha-1(tm329) II* | *elli-1; rha-1* | HCL226 crossed to SHG1941 | This study |

| | | | | |
|---|---|---|---|---|
| HCL190 | *npp-7(uoc21[mCherry::npp-7]) I* | *npp-7* | mCherry knockin by CRISPR-Cas9 to NPP-7 endogenous locus | JS Brown et al. 2023 |
| HCL214 | *rha-1(tm329) II; csr-1(uoc[gfp::csr-1]) IV* | *rha-1; csr-1* | HCL226 crossed to HCL105 | This study |
| HCL333 | *npp-7(uoc21[mCherry::npp-7]) I; rha-1(uoc24[gfp::rha-1]) II* | *npp-7; rha-1* | HCL207 crossed to HCL190 | This study |
| YY1492 | *mut-16(cmp3[mut-16::gfp::flag + loxP] I; znfx-1(gg634[HA::tagRFP::znfx-1]) II; pgl-1(gg640[pgl-1::3xflag::mCardinal]) IV* | *mut-16; znfx-1; pgl-1* | | CGC |
| HCL336 | *pgl-1(gg640[pgl-1::3xflag::mCardinal]) IV* | *pgl-1* | Outcrossed from YY1492 | This study |
| HCL334 | *rha-1(uoc24[gfp::rha-1]) II; pgl-1(gg640[pgl-1::3xflag::mCardinal]) IV* | *rha-1; pgl-1* | HCL207 crossed to pgl-1(gg640[pgl-1::3xflag::mCardinal]) originally from YY1492 | This study |
| HCL335 | *rha-1(uoc24[gfp::rha-1]) II; csr-1(gc029[degron::mCherry::3xflag::ha::csr-1]) IV* | *rha-1; csr-1* | HCL207 crossed to MHE69 | This study |
| HCL155 | *ego-1(uoc34[3xflag::gfp::ego-1]) I* | *ego-1* | 3xflag::GFP knockin by CRISPR-Cas9 to EGO-1 | This study |
| HCL337 | *ego-1(uoc34[3xflag::gfp::ego-1]) I; rha-1(tm329) II* | *ego-1; rha-1* | HCL226 crossed to HCL155 | This study |
| HCL338 | *rha-1(tm329) II; pgl-1(gg640[pgl-1::3xflag::mCardinal]) IV* | *rha-1; pgl-1* | HCL226 crossed to HCL366 | This study |
| HCL339 | *ego-1(uoc34[3xflag::gfp::ego-1]) I; pgl-1(gg640[pgl-1::3xflag::mCardinal]) IV* | *ego-1; pgl-1* | HCL155 crossed to pgl-1(gg640[pgl-1::3xflag::mCardinal]) originally from YY1492 | This study |
| HCL388 | *ego-1(uoc34[3xflag::gfp::ego-1]) I; rha-1(tm329) II; pgl-1(gg640[pgl-1::3xflag::mCardinal]) IV* | *ego-1; rha-1; pgl-1* | HCL339 crossed to HCL337 | This study |
| HCL340 | *ego-1(uoc34[3xflag::gfp::ego-1]) I; csr-1(gc029[degron::mCherry::3xflag::ha::csr-1]) IV* | *ego-1; csr-1* | HCL155 crossed to MHE69 | This study |
| HCL341 | *rha-1(uoc24[gfp::rha-1]) II; csr-1(gc029[degron::mCherry::3xflag::ha::csr-1]) IV; elli-1(uoc35) IV* | *rha-1; csr-1; elli-1* | CRISPR-Cas9 *elli-1* deletion strain made in HCL335 | This study |

| | | | | |
|---|---|---|---|---|
| HCL379 | *ego-1(uoc34[3xflag::gfp::ego-1]) I; rha-1(tm329) II; csr-1(gc029[degron::mCherry::3xflag::ha::csr-1]) IV* | *ego-1; rha-1; csr-1* | HCL226 crossed to HCL340 | This study |
| HCL376 | *wago-1(tor113[GFP::3xFLAG::wago-1]) I; csr-1(gc029[degron::mCherry::3xflag::ha::csr-1]) IV* | *wago-1; csr-1* | JMC219 crossed to MHE69 | This study |
| HCL389 | *wago-1(tor113[GFP::3xFLAG::wago-1]) I; rha-1(tm329) II; csr-1(gc029[degron::mCherry::3xflag::ha::csr-1]) IV* | *wago-1; rha-1; csr-1* | HCL376 crossed to HCL378 | This study |
| GKC1 | *mip-1(uae1) III* | *mip-1* | | CGC |
| HCL381 | *rha-1(uoc24[gfp::rha-1]) II; mip-1(uae1) III; pgl-1(gg640[pgl-1::3xflag::mCardinal]) IV* | *rha-1; mip-1; pgl-1* | HCL334 crossed to GKC1 | This study |
| CB4108 | *fog-2(q71) V* | *fog-2* | | CGC |
| HCL212 | *rha-1(tm329) II; fog-2(q71) V* | *rha-1; fog-2* | HCL226 crossed to CB4108 | This study |
| MHE69 | *ieSi64 [gld-1p::TIR1::mRuby::gld-1 3'UTR + Cbr-unc-119(+)] II; csr-1(gc029[degron::mCherry::3xflag::ha::csr-1]) IV* | *csr-1* | Degron knock in by CRISPR-Cas9 in MHE54 and cross with CA1352 (CGC). | P Quarato et al. 2020 |
| MHE113 | *ieSi64 [gld-1p::TIR1::mRuby::gld-1 3'UTR + Cbr-unc-119(+)] II; csr-1(gc029[degron::mCherry::3xflag::ha::csr-1]) ; gcSi1 [mex-5p::csr1[D769A]::GFP::tbb-2 3'UTR + Cbr-unc-119(+)] IV* | *csr-1* | Single copy insertion of mex-5p::csr1[D769A]::GFP::tbb-2 3'UTR in chromosome IV generated by MosSCI on strain EG6703 (CGC). Resulting MosSCI strain crossed with MHE69 to generate MHE113. | P Quarato et al. 2020 |
| MHE114 | *ieSi64 [gld-1p::TIR1::mRuby::gld-1 3'UTR + Cbr-unc-119(+)] II; csr-1(gc029[degron::mCherry::3xflag::ha::csr-1]) ; gcSi1 [mex-5p ::csr1::GFP::tbb-2 3'UTR + Cbr-unc-119(+)] IV* | *csr-1* | Single copy insertion of mex-5p::csr1::GFP::tbb-2 3'UTR in chromosome IV generated by MosSCI on strain EG6703 (CGC). Resulting MosSCI strain crossed with MHE69 to generate MHE114. | P Quarato et al. 2020 |
